# Characterizing the innate and adaptive responses of immunized mice to *Bordetella pertussis* infection using *in vivo* imaging and transcriptomic analysis

**DOI:** 10.1101/674408

**Authors:** Dylan T. Boehm, Melinda E. Varney, Ting Y. Wong, Evan S. Nowak, Emel Sen-Kilic, Jesse Hall, Shelby D. Bradford, Katherine DeRoos, Justin Bevere, Matthew Epperly, Jennifer A. Maynard, Erik L. Hewlett, Mariette Barbier, F. Heath Damron

## Abstract

*Bordetella pertussis* (*B. pertussis*) is the causative agent of pertussis (whooping cough). Since the 1990s, pertussis has re-emerged in the United States despite an estimated 95% vaccine coverage. Our goal was to characterize neutrophil responses and gene expression profiles of murine lungs in the context of vaccination and *B. pertussis* challenge. We utilized a bioluminescent neutrophil mouse model (NECre luc) to track neutrophil recruitment. NECre luc mice were immunized with whole cell vaccine (WCV), acellular vaccine (ACV), or a truncated adenylate cyclase toxoid (RTX) vaccine. Neutrophil recruitment was measured in live mice across time and corroborated by flow cytometry and other data. WCV immunized mice showed signs of neutrophilia in response to *B. pertussis* challenge. Mice immunized with either ACV or WCV cleared the challenge infection; however immunization with RTX alone was not protective. RNA sequencing revealed distinctive gene expression profiles for each immunization group. We observed an increase in expression of genes associated with responses to infection, and changes in expression of distinct genes in each vaccine group, providing a complex view of the immune response to *B. pertussis* infection in mice. This study suggests that combination of immunological analysis with transcriptomic profiling can facilitate discovery of pre-clinical correlates of protection for vaccine development.

## Introduction

Pertussis is a human disease primarily caused by a respiratory infection of the Gram-negative pathogen *Bordetella pertussis* (*B. pertussis*). The hallmark of pertussis is a distinctive whooping cough. What is surprising about pertussis is that the cause of the cough has never been elucidated, which highlights the fact that there are many under-researched aspects of this disease. Aerosolized *B. pertussis* bacterium are inhaled and adhere to airway respiratory epithelial cells through bacterial adhesins such as filamentous hemagglutinin (FHA), fimbriae and pertactin^1^. After colonization *B. pertussis* express multiple toxins including pertussis toxin (PT) and adenylate cyclase toxin (ACT). *B. pertussis* releases PT, which dysregulates the immune response through ADP-ribosylation of the G-protein α-subunit of cytokine receptors present on a range of leukocytes^2–5^. The secretion of PT has long range effects, ACT is thought to act locally on host cells by converting ATP into supraphysiological levels of cAMP, further dysregulating the host immune response^6^.

In the 1940s, an effective whole cell vaccine (WCV) was developed and as a result, basic research efforts on *B. pertussis* decreased. The WCVs were highly reactogenic and caused prolonged and unusual crying after administration, hyporesponsivness, and febrile convulsions ^7–9^. These issues led to the development of acellular vaccines (ACV), known today as DTaP/Tdap (hereafter referred to as ACV). The ACVs utilize an alum adjuvant and induce a Th2 response to several key virulence factors such as PT, FHA, fimbriae, and pertactin depending on vaccine formulations. Whereas the ACV induces a Th2 response in human and mice, WCV immunization and *B. pertussis* infection promote a Th1/Th17-type response^10–14^. Since the replacement of the WCV with the ACVs there has been a remarkable increase in the number of cases in the US, with the number of reported cases in 2012 equaling that of 1954. Multiple factors potentially played a role in the return of pertussis in the US. It has been documented that the Th2 response induced by ACV immunization may be inferior to the Th1/Th17 response directed by the WCV^15^. Additionally, ACV protection wanes significantly as early as one year after vaccination, and by 4 years after vaccination vaccine effectiveness is only 9%^16^. Furthermore, in a non-human primate model it has been shown that ACV immunized baboons challenged with *B. pertussis* are capable of carrying the infection and transmitting aerosolized *B. pertussis* to naïve baboons^17^. Additionally, the findings of an epidemiological study suggest that asymptomatic transmission of pertussis is indeed occurring in the human population, and has a role in the increase of pertussis incidents^18^. These recent findings demonstrate the shortcomings of the ACV. Utilizing new technologies that were not available during the developments of the WCV or ACVs we can further investigate differences between WCV and ACV protection. Adenylate cyclase toxin (ACT) is an essential virulence factor of *B. pertussis* and a known protective antigen^19–21^. ACT contains two main domains: adenylate cyclase and the Repeats-in-Toxin (RTX). In the absence of the AC domain, the protein is non-toxic and referred to simply as RTX^22^. In a previous study, sera from mice immunized with RTX contained antibodies capable of neutralizing ACT. ACT antigens are not included in any commercial formulations of ACVs and many have proposed that its inclusion could improve protection^6^.

Neutrophils play a role in both the innate and adaptive arms of the immune system. Following *B. pertussis* infection of a naïve mouse, neutrophils reach peak recruitment between 5 and 7 days after challenge^23,24^. This occurs after the bacterial burden has begun to decrease. Subsequent studies have determined that neutrophil depletion prior to *B. pertussis* infection in naïve mice does not increase bacterial burden. However, when immunized mice are challenged with *B. pertussis* following depletion of neutrophils the *B. pertussis* bacterial burden increase, demonstrating a protective role of neutrophils in immunized mice^25^. As previously mentioned, *B. pertussis* toxins PT and ACT both affect neutrophils and their recruitment^6,26–28^. These toxins contribute to a significant rise in the number of circulating white blood cells and in severe cases leukocytosis is linked to fatality^1^. Leukocytosis was first described in pertussis patients as early as the 1890s^29,30^. In the 1960’s, Stephen Morse observed leukocytosis following WCV immunization^31^, which he hypothesized was caused by PT, then known as leukocytosis-inducing factor^32^. Therefore, it can be conceived that the toxin associated effects on neutrophils would be neutralized following ideal *B. pertussis* immunization, and this raises the question of where and when neutrophils are recruited in an ACV or WCV protected mouse?

In this study, we utilized *in vivo* imaging systems (IVIS) to characterize the neutrophil responses to *B. pertussis* challenge in mice vaccinated with ACV or WCV. We implemented NECre luc mice, a luminescent neutrophil reporter mouse strain, previously used in a *Bacillus anthracis* infection model to track neutrophil recruitment following *B. pertussis* challenge^33^. To do so, we: 1) validated the NECre luc mouse as a model for tracking neutrophil recruitment in response to *B. pertussis* infection, 2) tracked the spatiotemporal localization of neutrophils in response to *B. pertussis* challenge in ACV and WCV vaccinated mice, and 3) and characterized vaccine-associated gene expression profiles of the infected lungs from each group using RNA sequencing. The NeCRE luc mouse^33^ was used to measure the presence and relative quantities of neutrophils in live mice during *B. pertussis* challenge. We then corroborated our findings with flow cytometry analysis to quantify neutrophils in the blood and airway of naïve and immunized mice. WCV immunization resulted in robust neutrophil recruitment and clearance of *B. pertussis,* but also resulted in morbidity of the NeCre luc mice. RNAseq analysis was performed on the lungs of naïve or immunized mice at both early and late time-points after challenge with *B. pertussis*. We observed specific gene signatures for each vaccine, and we also identified gene expression profiles that had not been associated with *B. pertussis* or immunization. Neutrophil specific gene expression corroborated our cellular analysis of the respiratory tract demonstrating increased expression of neutrophil specific genes in WCV immunized mice. Furthermore, we determined the T-helper cell gene expression profiles of lung transcriptomes and corroborated the activation of Th1 specific gene expression to increased levels of Th1-associated cytokine production in the lungs of WCV immunized mice. Utilizing the depth of data generated through RNAseq, we could classify specific B cell clonotypes present in the lung of each immunization group. This technique allowed for analysis of the diversity and frequency of the immunoglobulin repertoire of immunized groups. We hypothesize that these approaches can continue to be applied to other pathogen to host interactions to characterize the underpinnings to disease progression and immunological responses.

## Results

### ACV and WCV immunization of NECre luc mice results in clearance of *B. pertussis*, however WCV immunized mice experience increased morbidity and mortality

To characterize the effects of vaccination on neutrophil recruitment, NECre luc mice were immunized with the vaccines as indicated in Supplementary Table S1, and then boosted with the same vaccine 21 days later by intraperitoneal injection. We then compared the neutrophil recruitment in naïve, ACV, WCV, and RTX-only vaccinated NECre luc mice during *B. pertussis* challenge. At day 35 post initial immunization, NECre luc mic were challenged with virulent *B. pertussis* strain UT25 (Fig. 1) by intranasal instillation. A challenge dose of 2 × 10^7^ CFUs *B. pertussis* does not typically cause morbidity in immunocompetent mice such as CD-1 or C57B6/J (data not shown). We originally planned to euthanize 4 mice per group on each experimental day (1, 2, 4, and 9). However, 2 of 13 naïve challenged NECre luc mice were euthanized at day 2 due to morbidity (Fig. 2). At day 6, we observed additional unexpectedly morbidity in the WCV group, which required euthanasia of all remaining WCV immunized NECre luc mice (Fig. 2). Conversely, no morbidity was observed in the ACV and RTX immunized mice 9 days post *B. pertussis* challenge (Fig. 2). We were surprised to observe morbidity in the WCV immunized mice. Upon determining viable bacterial burden from lung, trachea, and nasal lavage, we observed clearance of *B. pertussis* to our limits of detection in both ACV and WCV immunized mice compared to naïve infection (Fig. 3abc). Taken together, these data suggested that morbidity of the WCV immunized mice was not fully attributed to bacterial burden. Additionally, we determined that vaccination with RTX resulted in 100 percent survival out to 9 days post-challenge (pc), while bacterial burden were similar to naïve infected mice, suggesting that RTX immunization did not improve bacterial clearance (Fig. 3abc).

**Figure 1:**
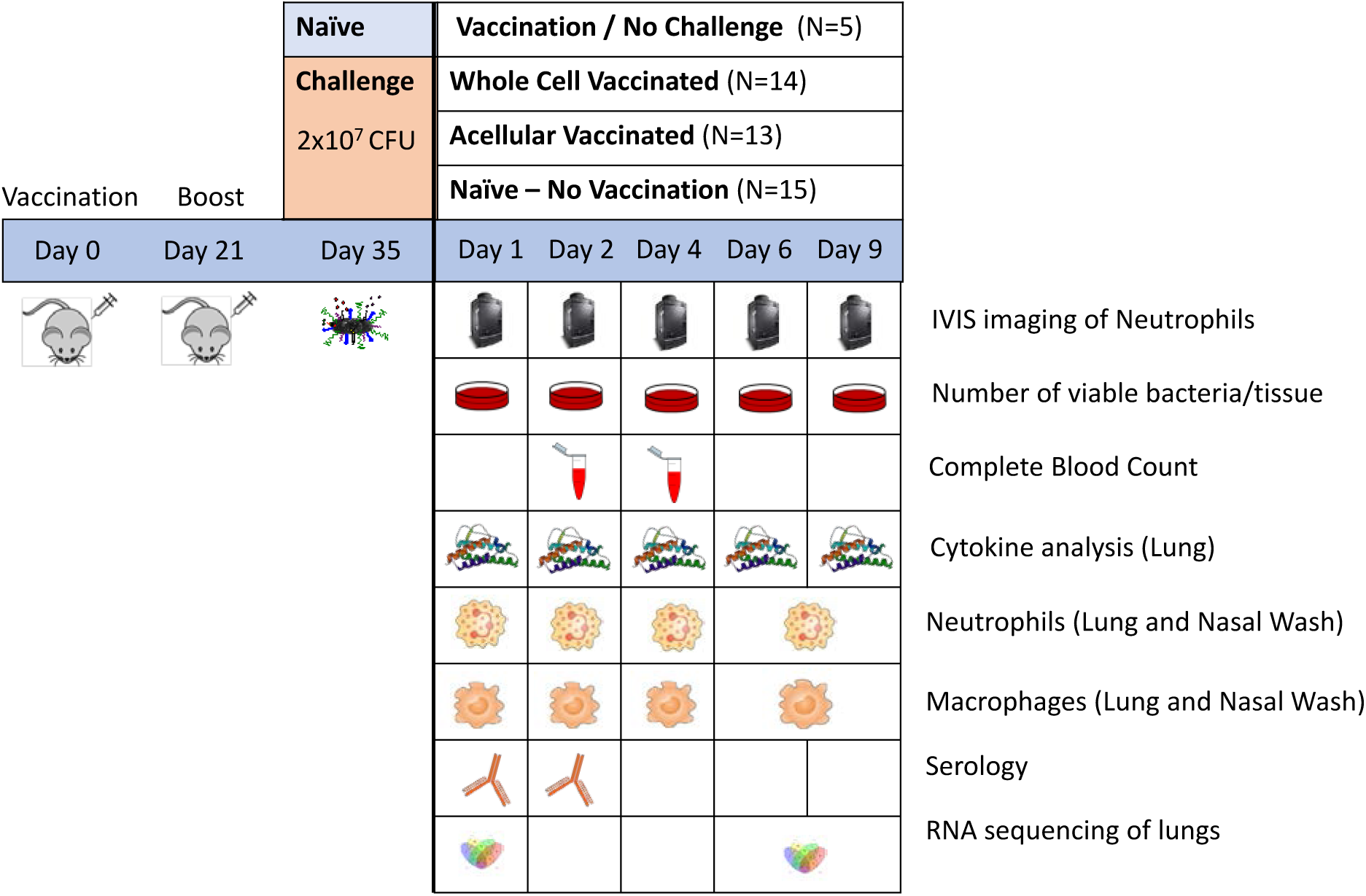
Experimental workflow for vaccination, *B. pertussis* challenge, sample acquisition, and analysis of this study. Schematic diagram of experimental design showing vaccination schedule, immunization groups, *B. pertussis* challenge, euthanasia, sampling, and analysis method on days 1,2,4, 6, and 9 days pc in NECre luc mouse model. Mice were initially vaccinated and then given a booster dose of the same vaccine at day 21. A challenge dose of 2 × 10^7^ viable bacteria was administered by intranasal inhalation at day 35. IVIS imaging of neutrophils was monitored throughout study. The overall numbers of mice per group are indicated. At time-points shown bacterial burden, complete blood cell counts, lung cytokine profiles, lung and nasal wash neutrophil cells, serum antibody titers, and lung transcriptome were determined. To characterize the effects of vaccination, the vaccinated groups were compared to a group of not vaccinated and not challenged controls.

**Figure 2:**
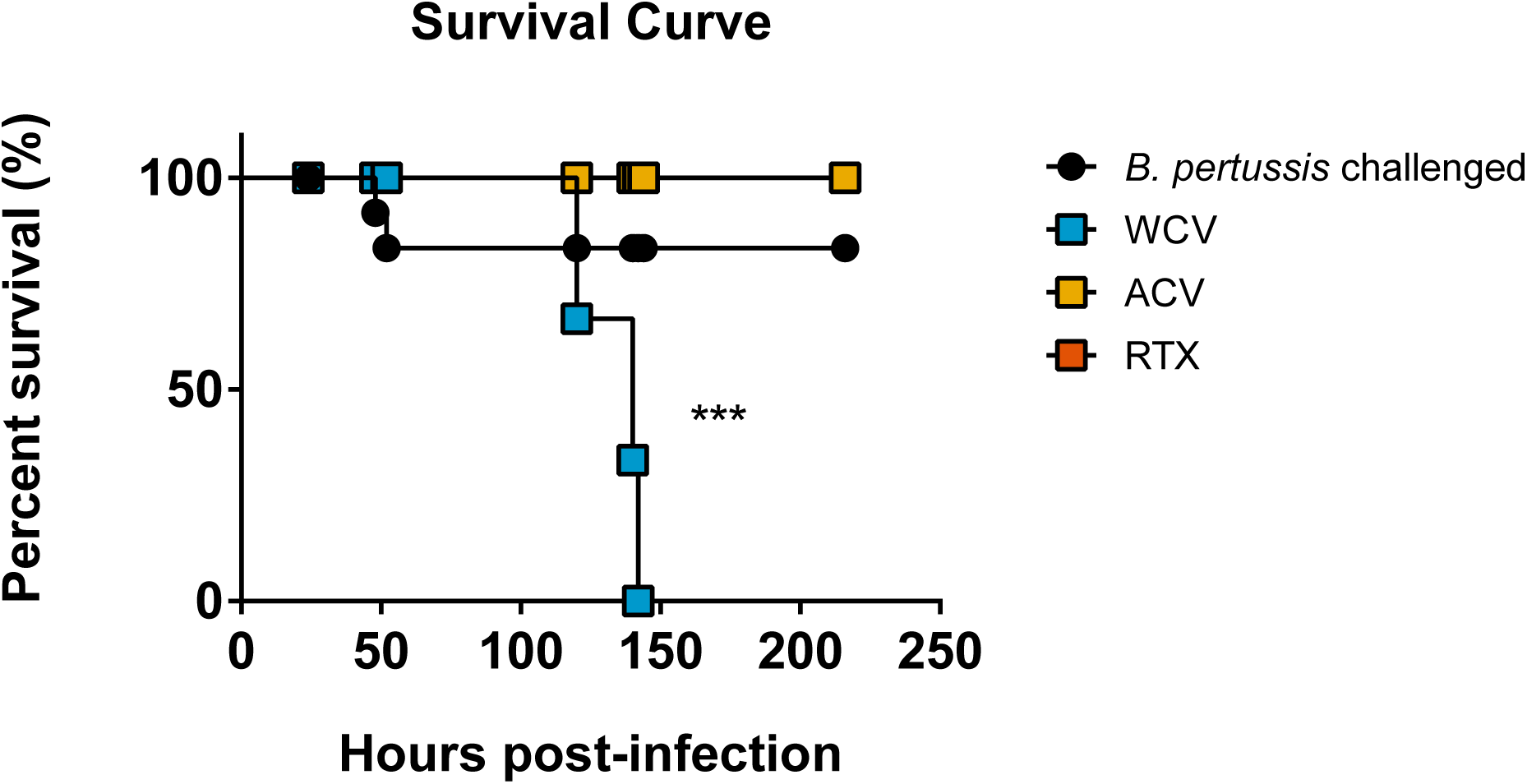
Kaplan-Meier survival curves of NECre luc mice according to immunization group. The survival percentage of remaining NECre luc mice (prior to scheduled euthanasia) immunized with WCV, ACV, RTX, and *B. pertussis* challenged, or not vaccinated and challenged with *B. pertussis*. Log-rank (Mantel-Cox) test: \*\*\**p< 0.0005.* In Fig. 1 the time-points where mice were euthanized for analysis are indicated. In this survival curve, we are only showing the mice became morbid throughout the study timeframe.

### Analysis of serological responses of immunized mice

As expected, the ACV immunized mice were protected against challenge as they rapidly cleared the *B. pertussis* challenge dose (Fig. 3abc). WCV immunized mice cleared *B. pertussis,* but this response was delayed compared to the ACV mice. To determine the efficacy of immunization on antibody production, we performed serology analysis. PT is a common antigen of both the ACV and WCV and we observed a significantly higher concentration of anti-PT in the ACV mice compared to the WCV (Fig. 3d). Immunization of NECre mice with RTX with alum adjuvant resulted in high anti-RTX antibody titers (Fig. 3e) despite the fact that we did not see increased clearance due to RTX immunization. In another study, we immunized CD1 mice with RTX and alum adjuvant and we also saw no clearance of *B. pertussis* compared to non-vaccinated mice (data not shown). Our data suggest that, while immunization with RTX induces antibody production, it is not sufficient to protect mice from *B. pertussis* infection as a single, antigen vaccine.

**Figure 3:**
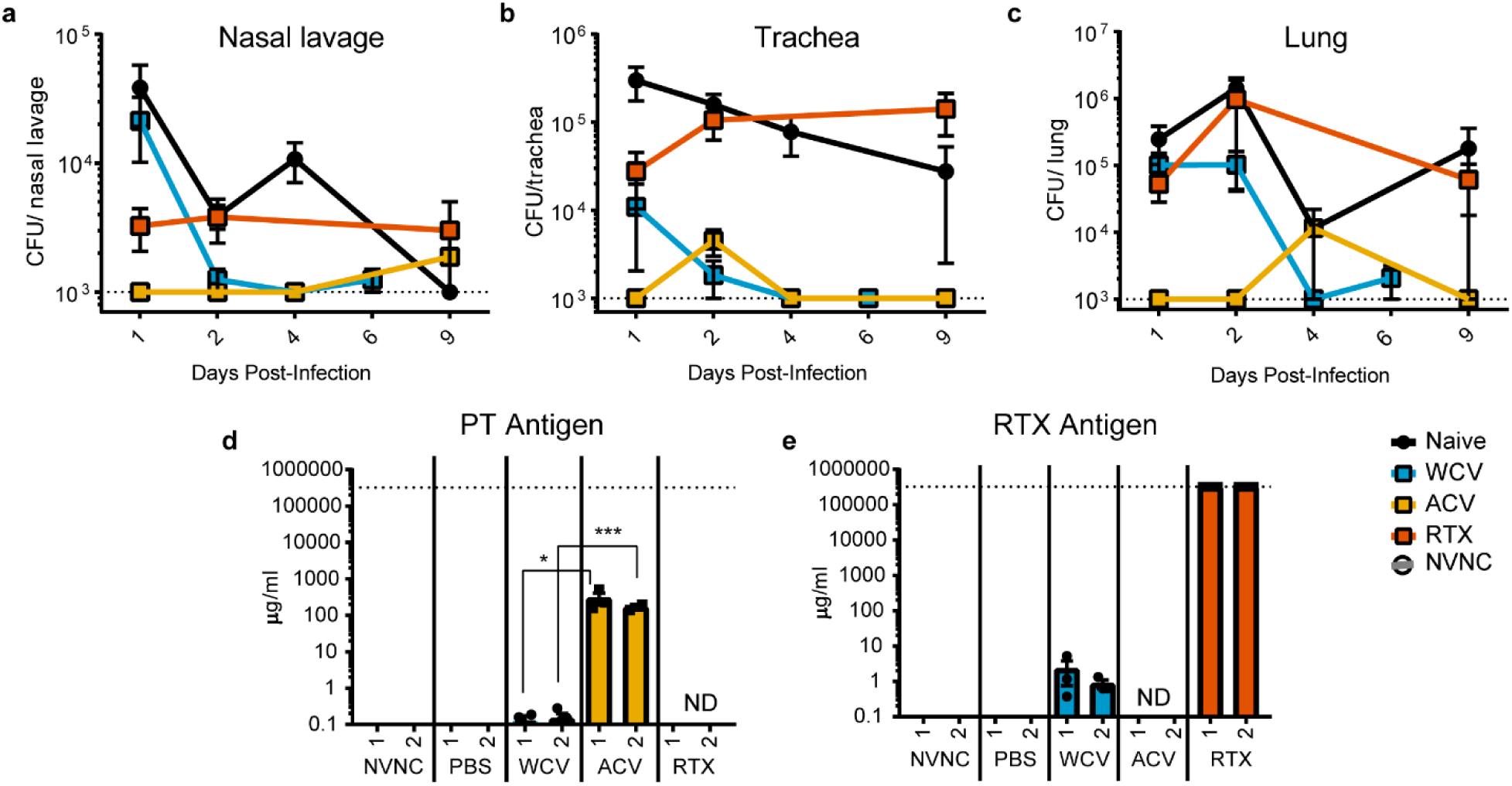
Bacterial burden in respiratory tissue and serological responses to *B. pertussis* challenge of immunized and naïve NECre luc mice. Mice were vaccinated with PBS control vehicle, WCV, ACV, or RTX then *B. pertussis* challenged. At days 1,2,4,6 and 9 pc the bacterial burdens were determined by culturing of (A) nasal lavage, homogenates of (B) trachea and (C) lung on BG agar. The dashed line at 1000 CFUs represents the lower limit of detection, due to plating. Data in each group were compared to PBS control using an unpaired two-tailed t-test. Significant differences are not indicated on the graphs for clarity (Supplementary Table 2 indicates all statistics performed).

### Spatiotemporal IVIS imaging of neutrophils in naïve and vaccinated NECre luc mice

Neutrophil accumulation in NECre luc mice were visualized on days 1, 2, 4, 6 and 9 post-challenge. Mice were IP injected with the high sensitivity luciferin CycLuc1^33^ before imaging. A Lumina II IVIS (Xenogen) was used to capture luciferase-driven neutrophil luminescence at indicated time-points (Fig. 4 and Supplementary Fig. 1). Luminescent signal of the whole body (Fig. 4d) or nasal cavity (Fig. 4e) of vaccinated mice was then compared to signal from naïve, non-infected mice and calculated as fold change. At day 1 post-challenge, we detected significantly higher luminescence in naïve challenged mice compared to naïve non-challenged mice, suggesting that there is a higher neutrophil accumulation in challenged mice. Furthermore, whole body luminescence in WCV immunized and challenged mice was significantly higher than the whole body luminescence of naïve, ACV and RTX *B. pertussis* challenged mice at day 1 (Fig. 4d). In general, luminescence signals of all groups decreased across days 2 and 4. However, in the WCV group at day 6, we detected an increase in signal in the nasal cavity (71-fold) and the whole body (47-fold) compared to naïve not challenged mice (Fig. 4de). Due to variability between animals, this increase was not significant compared to other groups. At this point WCV immunized mice were morbid and required euthanasia. Taken together this data suggests that WCV immunization prior to challenge resulted in increased neutrophil accumulation compared to other vaccination groups. We next aimed to corroborate our IVIS luminescence data with flow cytometry analysis of respiratory tissues.

**Figure 4:**
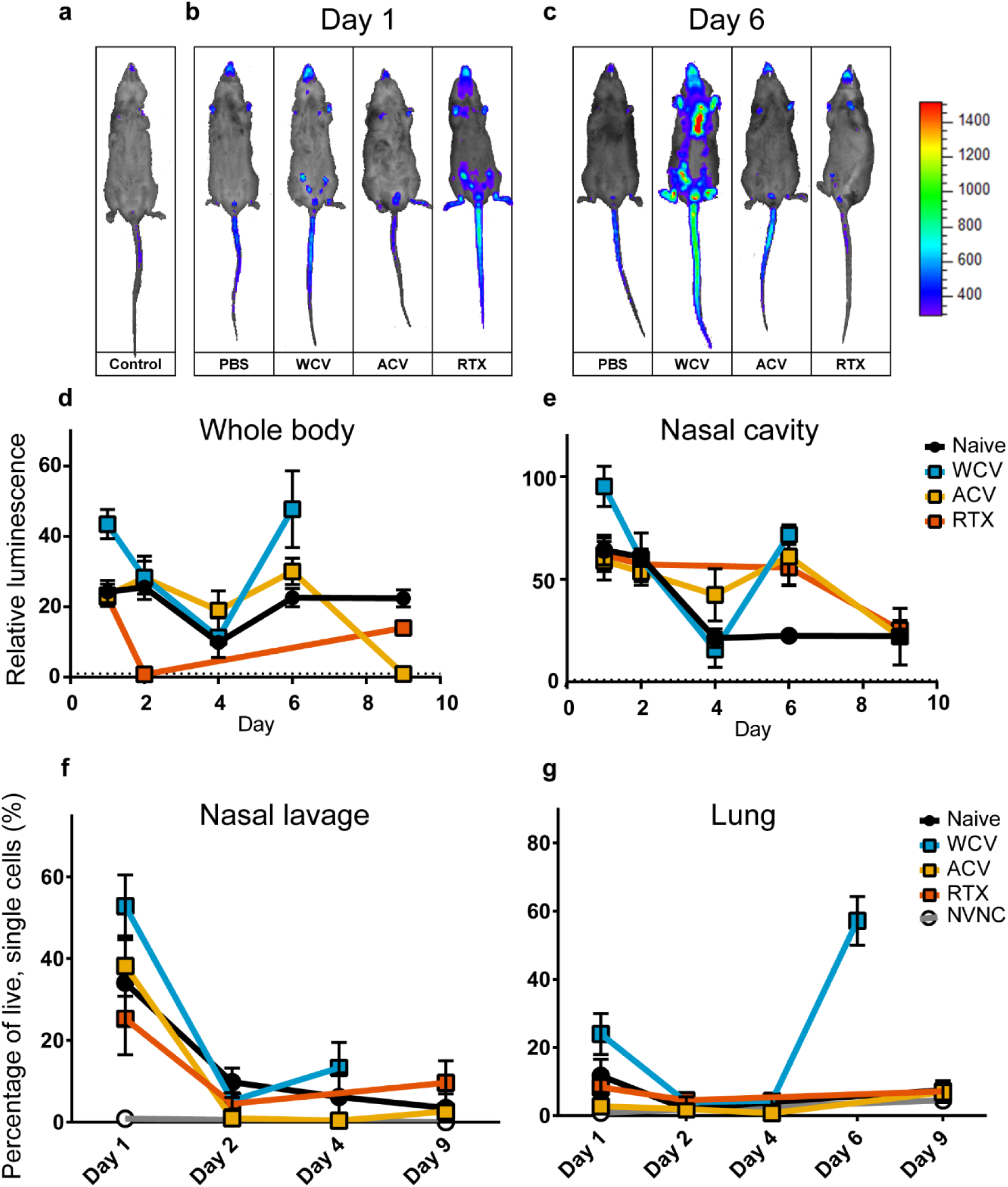
IVIS imaging and flow cytometric analysis of naïve and immunized NECre luc mice post challenge with *B. pertussis.* IVIS imaging of luminescent neutrophils in anesthetized mice following luciferin injection. (A) Representative images of naïve, not *B. pertussis* challenged NECre luc mice. (B) Representative images of PBS control, WCV, ACV, and RTX immunized and *B. pertussis* challenged NECre luc mice at day 1 pc and (C) day 6 pc (D and E). Luminescence was measured on Xenogen IVIS Lumina II. Relative luminescence levels quantified by fold change of emitted photons/second of NECre luc mice following *B. pertussis* challenge (N=3-5) compared to average luminescence of naïve, not challenged NECre luc mice (N=5). Neutrophil luminescence of (D) whole animal signal and (E) nasal cavity was determined at days 1,2,4,6, and 9 pc. (f) Quantification of the percentage of live, single cells classified as neutrophils (GR-1+CD11b+) detected in nasal lavage (f) and lung homogenate (g). Significant differences are not indicated on the graphs for clarity, but are included in Supplementary Table 3.

### Validation of neutrophil infiltration in respiratory tissue by flow cytometry

Luminescent signal of luciferase-expressing cells correlates with the localization of neutrophils in NECre luc mice^33^. To confirm this, we measured the relative number of neutrophils by performing flow cytometry analysis on respiratory tissue following IVIS imaging. Single cell suspensions from lung tissue or nasal lavage were labelled with antibodies recognizing cell surface markers. We defined neutrophils as CD11b+GR-1+ live cells (Supplementary Fig. 2). Similar to the IVIS data, all *B. pertussis* challenged groups had significantly higher neutrophil percentages in nasal lavage compared to naïve non-challenged mice (Fig. 4f). At day 1 post *B. pertussis* challenge, WCV immunized mice had a significantly higher percentage of neutrophils than ACV immunized mice in nasal lavage. The percentages of neutrophils decreased from day 1 to day 2 in all vaccinated groups, as well as the naïve group. At day 4, WCV immunized mice exhibited higher levels of neutrophils than those observed at day 2 (Fig. 4g). Correlating with the nasal lavage, WCV mice exhibited significantly higher percentages of neutrophils in the lungs than ACV mice. Lung neutrophils at day 4 were not statistically significant between any of the groups. However, by day 6 there was a 53% increase in lung neutrophils compared to day 4 in the WCV group, which directly correlates with the increased luminescence signals measured by IVIS imaging (Fig. 4g).

### WCV immunized NECre luc mice induce a more robust Th1 immune response compared to ACV immunized mice

It has been previously documented that upon *B. pertussis* challenge, WCV immunized mice elicit a Th1/Th17 immune response, similar to a naïve infection while ACV immunized mice generate primarily a Th2 response^10–12,34^. We hypothesized that similar responses are triggered by WCV and ACV in the NECre luc model. To test this hypothesis, we determined the cytokine profiles in lungs of immunized and challenged mice using electrochemiluminescent sandwich immunoassays. The Th1 response was determined by increased levels of IFN-γ, IL-12p70, IL-1β and TNF-α (Fig. 5ab), while the Th17 response was qualified by increased IL-17A and IL-6 (Fig. 5cde). Th2 responses were associated to increased levels in IL-4 and IL-5.

As expected, we observed an increased Th1 response in the WCV vaccinated NECre luc mice compared to the ACV group. We also observed an increase in the Th1 response in naive challenged mice compared to non-challenged mice or the ACV group. Interestingly, the highest levels of IL-17 were observed in WCV NECre luc mice (Fig. 5c). The levels of Th2-associated cytokines, IL-4 and IL5 were similar between the WCV and ACV groups. (Supplementary Fig. 3). The lung cytokine profile from NECre luc mice immunized with RTX resembled a profile of naïve *B. pertussis* challenged mice with the exception of IL-6, which was lower in the RTX group. Taken together these data from the NECre luc mouse model corroborates previous findings by displaying a strong Th1/Th17 response in response to *B. pertussis* in WCV vaccinated mice.

**Figure 5:**
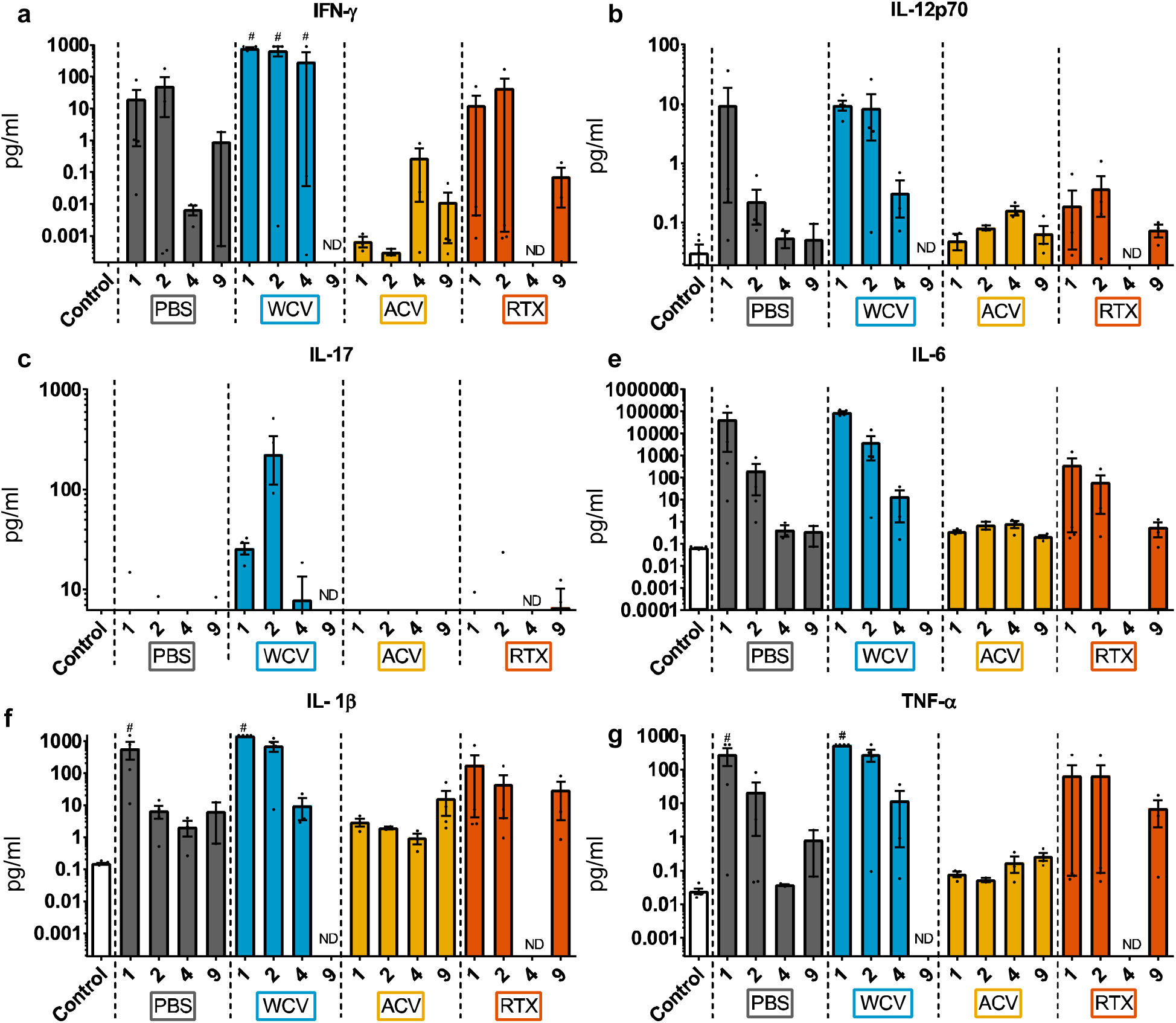
Analysis of cytokine profiles from lungs of naïve and immunized NECre luc mice post challenge with *B. pertussis.* Th1 associated cytokines from the supernatant of lung homogenates were analyzed at days 1,2,4 and 9 pc. Cytokines (a) IFN-γ, (b) IL-12p70, (c) IL-17, (d) IL-6, (e) IL-1β, (f) TNF-α were quantified using electrochemiluminescence immunoassays. Significant differences are not indicated on the graphs for clarity but, are included in Supplementary Table 4. ND: Sample not determine, #: data above upper limits of detection

### Characterizing the gene expression profiles of the lungs of immunized and naïve mice after *B. pertussis* challenge

Current and past *B. pertussis* immune response studies have focused on a limited set of known immunological responses to *B. pertussis* challenge such as: bacterial burden, antibody response, recruitment of phagocytes, cytokine profiles, and others ^35,36^. Whooping cough is a toxin-mediated disease, due mainly to the activity of pertussis and adenylate cyclase toxins. These toxins are released and affect a wide array of host cells^4,25^. It is therefore conceivable that *B. pertussis* infection would have a broader effect on the respiratory tissue beyond the induction of predictable immune response factors. To address this, we employed RNA sequencing (RNAseq) to observe the effect of *B. pertussis* challenge on the lungs of immunized and naïve NECre luc mice at a transcriptomic level. This allowed us to characterize the transcriptome response to infection and vaccine-induced responses. We hypothesized that regardless of the vaccine administered, we would identify a set of genes differentially regulated in all challenged mice. Additionally, we sought to discern the unique gene expression profiles in response to challenge in each immunized group.

To perform RNAseq analysis, lungs from NECre luc mice were harvested at an early time-point (day 1) following *B. pertussis* challenge and at a late time-point (day 6 or 9). RNA from the lungs of the mice was isolated. Libraries were prepared and sequenced on the Illumina HiSeq 1500 platform. To characterize transcriptional responses to *B. pertussis* challenge in naïve or vaccine mice (ACV, WCV, RTX), we compared the challenged samples to control mice that were not immunized nor challenged. Based on this, we expected to determine both the immunological (innate and adaptive) and the non-immunological responses to *B. pertussis* challenge. We determined the differentially expressed genes (DEGs) for both early (day 1) and late (day 6 or 9 depending on group) time-points. The total numbers of differentially regulated genes are shown in Fig. 6ac. Overall, there were more differentially genes at the later time-point than early after challenge (Fig. 6ac). At day 1 post-challenge, the WCV group had the most genes activated (Fig. 6a, Supplementary Fig. 5). Of the genes that were upregulated at day 1 in the WCV, 29.7% were not found upregulated in the other groups (464 of 1,560). At the late time-points, there were more common upregulated genes and only 14.1% of the genes were unique to the WCV. We found that 12.6% of unique upregulated genes (196) were common to all groups either challenged with *B. pertussis* following immunization or naïve challenge, suggesting that these genes were common to infection (Fig. 6b). At the later time-point, 19.8% of unique upregulated genes (867 of 6,595) were shared by the WCV, ACV, RTX groups, while this number was much lower at the early time point (8.5% of DEGs, 133 of 1,560) (Fig. 6bd). Cumulatively, this data suggests that: 1) WCV vaccination induces a more diverse early response to *B. pertussis* challenge due to the higher number of unique genes compared to other groups, and 2) at the later time-point all vaccinated groups have a more similar lung transcriptome.

**Figure 6:**
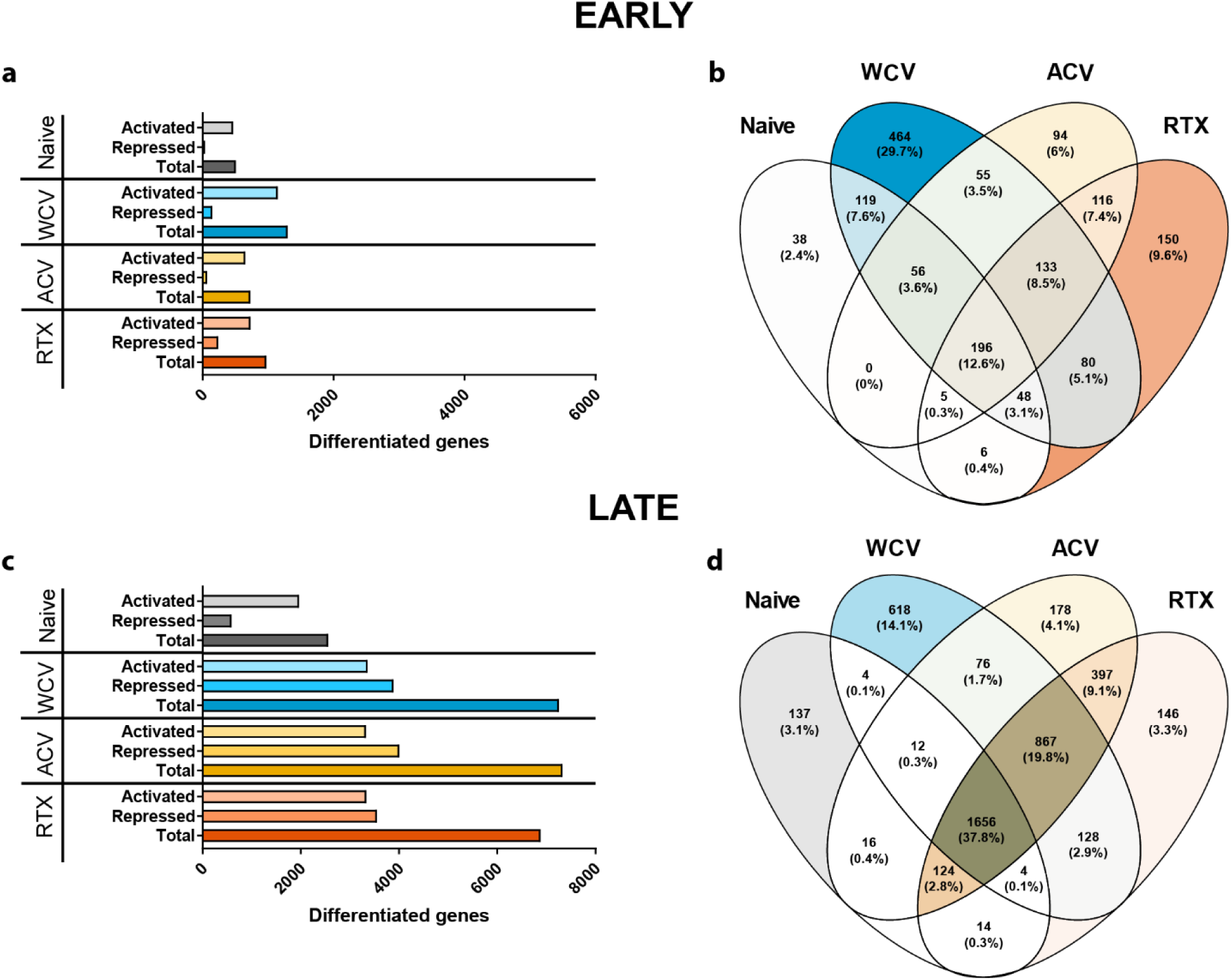
Lung transcriptome profile of vaccinated and challenged NECre luc mice. RNA sequencing was performed on total RNA from homogenized lung on days 1 (Early) or 6/9 (Late) following challenge with *B. pertussis.* (a) The total number of statically differentiated genes found in lung transcriptome at early (a) or late (c) time-points. Statistically differentiated genes were categorized as those that were activated or repressed. Venn diagram of statistically differentiated genes either unique or common to vaccinated or naïve groups at early (b) or late (d) time-points.

Ingenuity Pathway Analysis (IPA) was performed to determine enrichment analyses of gene expression signatures in the context of each immunization. Using IPA Knowledge-based enrichment analysis, statistically significant genes were grouped into disease and function categories. A system of genes was considered enriched based on Fisher’s exact test on the ratio of represented genes. At the early time-point we, observed that the largest changes were associated with genes involved in immune cell trafficking of the WCV group (Fig 7a). Genes with the highest fold changes within this system were associated with activation and migration of leukocytes (Fig. 7b). Additionally, we observed significant enrichment of genes associated with the humoral and cell-mediated immune responses (Fig. 7b). Pertussis toxin affects multiple aspects of the cardiovascular system such as blood pressure, leukocyte migration, etc^7,37,38^. In both naïve and immunized mice, we observed gene expression signatures related to cardiovascular diseases (Fig. 7ab). ACV immunization results in high amounts of anti-PT antibodies (Fig. 4d), and we observed fewer genes significantly altered in the ACV immunized mice, suggesting positive effects of anti-PT antibodies on neutralization of toxin activity. At the late time-point, immune cell trafficking gene expression seemed to decrease in naïve, ACV, and RTX groups; however, the expression of these genes was still high in the WCV group. As expected, immune-related gene signatures dominated the responses to *B. pertussis* infection. However, thousands of genes significantly dysregulated due to infection at both time-points were not related to immune response or were not sufficiently characterized to be assigned a function. Analysis of the lung transcriptomes corroborated the findings of flow cytometry and cytokine profile analyses, further supporting that there is an increase in leukocyte activity in the respiratory tissue of WCV immunized mice compared to ACV or RTX immunized mice following *B. pertussis* challenge.

**Figure 7:**
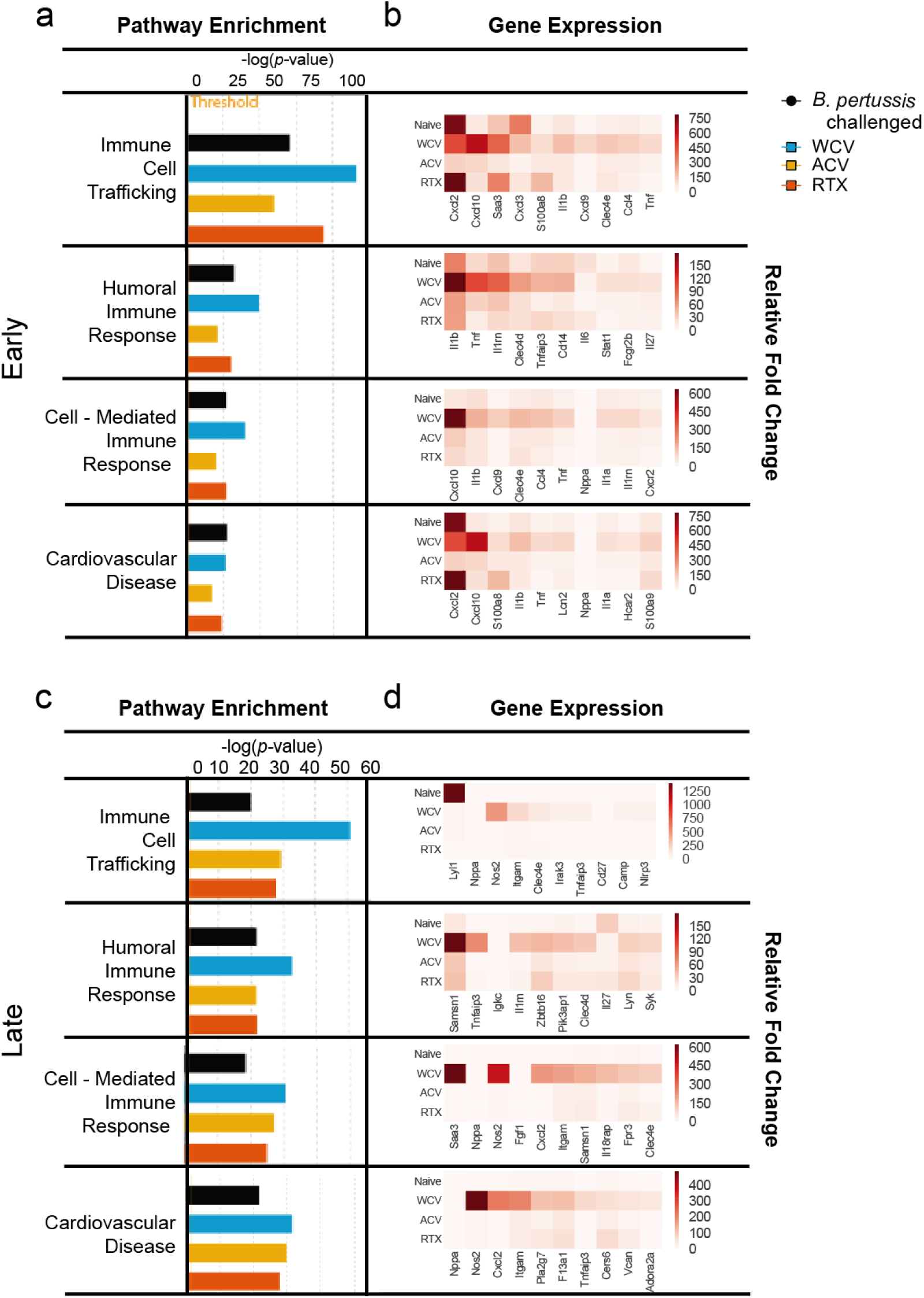
Enrichment analysis and relative fold-changes of genes upregulated following *B. pertussis* challenge. Early (day 1)(a), and late (day 9)(b) IPA comparative analysis of lung transcriptomes from vaccinated and non-vaccinated/challenged groups was performed, gene enrichment *p*-values are represented in –log scale. *P*-value was calculated by IPA software based on the number of genes found in a certain data set to the total number of genes associated to a particular function in the IPA knowledge base. Threshold indicates the significance (*p*<0.05, Fisher’s exact t-test) Black-PBS, Blue – WCV, Yellow-ACV, Red – RTX. Relative fold changes of ten highest genes associated with a particular function annotation present in either of the experimental groups.

### Analysis of neutrophil-specific gene signatures

Since we observed that neutrophils accumulate in the lung of WCV immunized NECre luc mice, we chose to delve further into the changes in expression of genes associated with neutrophil activity. To compare the neutrophil response between the groups, we selected genes with GO terms associated with neutrophil migration and activation and plotted the relative fold changes for this subset of genes for each immunization group (Fig. 8). The expression levels of these genes from either the naïve challenged, or immunized and challenged groups was compared to naïve, not infected control mice (Fig. 8).

**Fig. 8:**
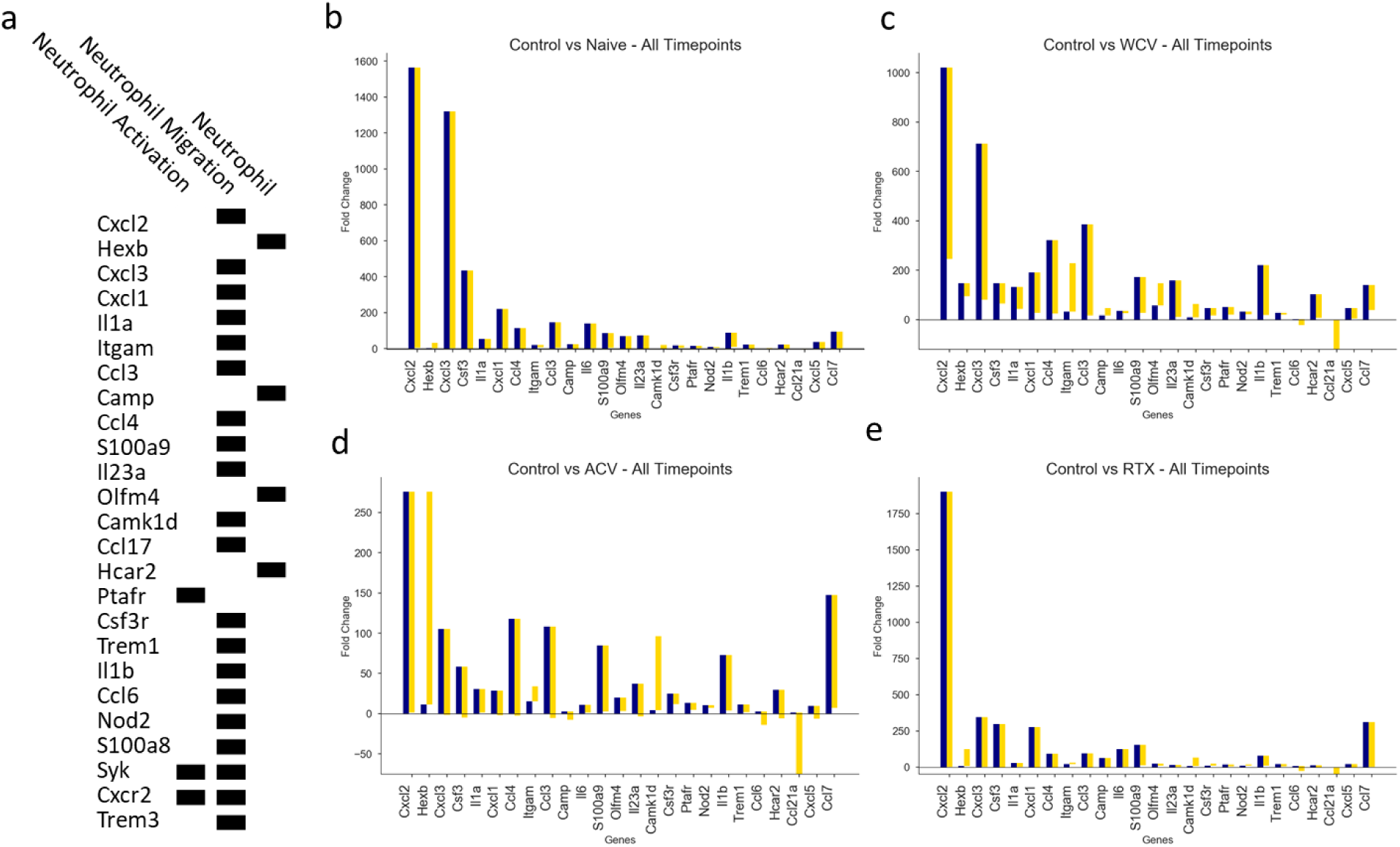
Differentiated genes associated with neutrophil recruitment at early and late time-points following *B. pertussis* challenge. Gene expression profiles are shown for (a) Naïve, (b) WCV, (c) ACV, and (d) RTX immunized and challenged mice. Blue and gold bars indicate the fold change compared to control non-challenge non-vaccinated mice at early and late time-points, respectively. Genes shown are the 25 genes with the highest variance across all groups between early and late time-point. Genes were ranked based on variance in fold change across groups for the early and late time-points, separately.

Cxcl2 is a chemokine produced by monocytes and neutrophils at sites of infection^39^. Previously it has been shown that convalescent mice previously challenged with *B. pertussis* express lower amounts of *cxcl2* when re-challenged with *B. pertussis* than naïve mice^40^. This suggests that the adaptive responses in these mice negate the need for the innate production of Cxcl2. Supporting this hypothesis, we observed the highest expression levels of *Cxcl2* in naïve and RTX immunized mice, which had similar bacterial burdens (Fig. 7bd and Fig. 8be). The induction of *Cxcl2* expression was lower in ACV vaccinated mice than naïve mice (Fig. 8ac). The expression of *Cxcl2* also increased in the WCV mice, but remained high at the late time-point (Fig. 8b). Our serological analysis indicated that WCV immunization only induced minimal anti-PT production, whereas ACV induced high amounts of anti-PT (Fig. 3d). PT has been shown to induce production of Cxcl2 in TY10 cerebral endothelial cells which further suggests a causal relationship between non-neutralized PT and Cxcl2 production^41^.

All of the groups showed increases in a number of other C-X-C and C-C family chemokines (*cxcl3, cxcl1, ccl3, ccl4, ccl6,* and *ccl17*). *Cxcl1* gene expression was increased at day 1 in all groups compared to the non-infected control (Fig. 8). We also observed an increase in *IL-23* expression in the WCV group (Fig. 7b), which might be associated with the IL-17 cytokine response measured in the WCV lung tissue. Tumor necrosis factor (*tnf*) gene expression was increased in WCV, RTX, and ACV mice compared to non-infected mice. However, *Tnf* expression levels decreased in ACV mice and were lower than those observed in uninfected mice at day 9 (Supplementary Fig. 4). Ross *et al.* demonstrated that IL-17A promotes production of CXCL1 during *B. pertussis* challenge^10^. In our cytokine analysis, only the WCV mice had sufficient IL-17 to be detected in the lung homogenates (Fig. 5c). Taken together these data suggest that although we did not observe high *IL-17A* gene expression in the transcriptome or in the IL-17 cytokine analysis, there could still be sufficient IL-17A in the Naïve, ACV, and RTX challenged mice that could account for the increased C*xcl1* gene expression.

We observed that *S100a9* and *S100a8* were highly upregulated in WCV and RTX groups (Fig. 7 and Fig. 8b). S100a9 is a major calcium-binding protein of neutrophils, which forms a heterodimer with S100a8, and promotes polymerization of microtubules. Interestingly, S100a8/a9 is produced mainly by neutrophils and high plasma levels of S100a8/a9 correlate with higher blood neutrophil blood levels in heathy human adults^42^. S100a8/a9 promotes transendothial phagocyte migration by promoting tubulin polymerization^43^. Additionally, monocytes isolated from human patients with cardiac injury have been found to be particularly responsive to S100a8/a9 and secrete increased amounts of TNF-α and IL-6^44^. S100a8/a9 can induce neutrophil chemotaxis and adherence, but PT can block this function of S100a8/a9^45^. These data suggest that PT induction of leukocytosis may be related to the role of S100a8/a9 in leukocyte recruitment. S100a8/a9 expression was lowest in the ACV mice which had the highest amount of serum anti-PT.

Overall the WCV mice had higher expression of more neutrophil genes than any of the other groups (Fig. 7). Expression of these neutrophil signatures did not return to uninfected levels. These observations corroborates IVIS (Fig. 4de) and flow cytometry analyses (Fig. 4fg). Neutrophil gene expression in ACV mice was dynamic and returned to levels lower than uninfected mice by day 9. ACV mice did not have detectable *B. pertussis* in their lungs at day 9, indicating that the initial challenge does had been cleared (Fig. 3). ACV and WCV signatures were the most dissimilar, while the RTX and naïve mice transcriptome profiles were the most similar (Fig. 7). This correlates with the fact that RTX mice still have detectable *B. pertussis* in their nares, trachea and lungs at day 9 at levels similar to those observed in non-vaccinated mice. These data confirm our neutrophil observations but also enhance our analysis of the neutrophil dynamic in the context of vaccination status and *B. pertussis* challenge.

### B cell gene expression signatures in relation to immunization status

It has been established that B cells are required for the clearance of *B. pertussis* in mice and that ACV immunization protects through neutralizing antibodies^46,47^. To further investigate antibody production between WCV and ACV immunized mice, we aimed to determine the diversity of B cell clone expansion between vaccinated and non-vaccinated challenged mice. Using MiXCR software, the same illumina sequenced reads mentioned above were analyzed specifically for immunoglobulin profiling^48^. Individual B cell clones were identified by B cell receptor sequence and clonal expansion was accessed by immunoglobulin diversity and frequency between the vaccinated and challenged, naïve challenged, and non-challenged control groups. At the early time-point day 1 post-challenge, immunoglobulin diversity and frequency of B cells depicted a close relation between the ACV and RTX vaccination groups, as also did WCV and naïve challenged mice (Supplementary Fig. 6a). This response shifted by the late time-point (days 6 or 9), when we observed that WCV and ACV groups were more similar. Using this approach, we were able to determine the individual variable and joining segments abundance, and the frequency of which a particular was found. Not surprisingly, we observed the highest degree of diversity at the late time-point. WCV and ACV were the most similar due to a higher number and abundance of clonotypes (Supplementary Fig. 6b). We focused our analysis on the diversity of the third complementarity-determining region (CDR3) of the heavy Ig chain, because this region has been shown to be necessary for antigen specificity^49^. The highest abundant CDR3 region sequence, NQHLFW, was present only in the WCV immunized mice, while the other 10 highest CDR3 regions were shared across vaccine groups at the early and late time-points (Supplementary Table 6 and Supplementary Table 7). Together this data suggests that clones generated from acellular based vaccines (ACT and RTX) are more closely related because they are generated to a limited number of antigens. However, by the late time-point, proliferation of specific B cell clones generated from WCV has created the same diversity seen in the ACV clones. The B cell clonotype analysis suggests that we can aim to further understand these populations in the future but we will need to increase our sequencing depth by directly isolating the cell populations from the immunized mice instead of analyzing the full lung tissue. It may also be interesting to probe the T cell receptor clonotypes using the same method.

### Discussion

In this study, we compared the innate and adaptive immune responses of immunized NECre luc mice vaccinated with ACV or WCV vaccines, to mice immunized with a truncated ACT toxin absorbed to alum (RTX), or non-vaccinated naïve mice following challenge with *B. pertussis.* ACV and WCV immunized mice cleared *B. pertussis* challenge but distinctive immune responses were observed. We also determined that the RTX antigen alone was not protective as a single antigen vaccine with alum adjuvant in NeCre luc mice. We utilized IVIS imaging to track the recruitment of neutrophils to the respiratory tract of challenged mice from days 1 to 9 post-challenge. The data obtained illustrated a dramatic increase in the amount of neutrophils recruited to the lungs and nasal cavity of WCV immunized mice compared to ACV immunized mice when followed by *B. pertussis* challenge. These data were also supported by flow cytometry and cytokine analysis. Furthermore, by performing RNA sequencing on the lung transcriptome during infection in vaccinated or naïve mice, we described unique gene expression profiles depending on the vaccination status of the mice. We also used RNA sequencing to begin to describe the immunoglobulin diversity induced by each vaccine.

Upon initial *B. pertussis* challenge, we observed activation of neutrophil associated genes in all groups, however it was only in the WCV immunized mice that we saw increased neutrophil genes expression still elevated at the late time-point. Neutrophilia and elevated neutrophil related genes expression at the late time-point, when bacterial numbers in the airways are greatly reduced, indicates either over activation of neutrophil “chemokine storm,” or possibly a lack of neutralization of bacterial toxins. Multiple studies suggest that PT is responsible for neutrophilia and leukocytosis following *B. pertussis* infection^13,32,50,51^. Immunization with the ACV induced 1,146- fold more anti-PT antibody production than WCV immunization at day 2 (Fig. 3d). It is possible that the low levels of anti-PT production in WCV mice resulted in insufficient neutralization of PT, resulting in exacerbated neutrophilia. Surprisingly, the highest neutrophil accumulation was determined at day 6 post-challenge when bacterial levels were similar to those of ACV protected mice (Fig. 3). These findings suggest that while WCV immunization is protective at clearing infection in NECre luc mice, it also induced severe neutrophilia. Interestingly, the neutrophil response in WCV mice decreased at days 2 and 4 but highly increased at day 6 (Fig. 4). At day 2, we observed a significant increase in IL-17 in the lungs of WCV mice (Fig. 5c) that could result from the increased IL-17 expression in these mice. Stephen Morse observed dose-dependent leukocytosis in mice following vaccination with killed *B. pertussis*^31^. In this study, NECre luc mice were immunized with 1/5^th^ the human dose of WCV. We now realize that this dose is likely well above the proportional weight of a mouse compared to a human. It is possible that this high vaccine dose induced the hyper leukocytosis, similar to the dose-dependent increase that Morse observed. It is known that hyperleukocytosis is associated with death in infant cases, and that PT is responsible for inducing leukocytosis^1,7,52^. Furthermore, mice and baboons infected with *B. pertussis* and then treated with anti-PT antibodies had lower levels of leukocytosis compared to non-infected controls^50^. An epidemiological study found that in unvaccinated individuals with pertussis, 72% of patients experienced leukocytosis^53^. In this study, we observed that WCV immunized NECre luc mice experienced neutrophilia and morbidity, which highlights the potential issues of using WCVs. It would be interesting to determine the relationship of neutrophilia and WCV immunization in epidemiological past studies but we have not been successful in identifying a study that specifically looked at neutrophilia because studies most note general leukocytosis.

In NECre luc mice, the ACV was clearly more protective and less detrimental to the mice than the WCV. Truncated and full-length ACT purified from *B. pertussis* culture has been shown to be a protective antigen^21^. Wang *et al.* described the RTX region of ACT as highly immunogenic and easily purified as a recombinant protein^22^. We have observed that immunization of CD1 mice with RTX and alum adjuvant results in high anti-RTX titers (data not shown). Here, we immunized NECreluc mice with RTX and alum, but no protection was observed (Fig. 3abc). The previous studies were performed with strain 18323 and the antigen was directly isolated from *B. pertussis*^21,54^. It is now known that strain 18323, is an outlier compared to most other global *B. pertussis* strains and Guiso *et al*.^21^ noted that strain 18323 produces less PT than the Tohama I type strain. From our work, we know that UT25 produces more PT and ACT than Tohama I (data not shown). It is possible that due to the increased PT levels produced by UT25, vaccination with RTX was not sufficient to block colonization and proliferation *in vivo*. Although RTX immunization did not result in clearance of *B. pertussis* from NECre luc mice, we did observe a reduction in the pro-inflammatory cytokine IL-6 in the RTX group compared to not vaccinated and WCV vaccinated and challenged mice (Fig. 5e). PT and ACT have both been shown to induce the production of IL-6 in human cell lines^55,56^. The anti-ACT antibodies generated by RTX vaccination may play a role in reducing the levels of IL-6 due to reducing the activity of ACT. We hypothesize that it is necessary to neutralize both PT and ACT to provide optimal protection and in future studies, we will test RTX as an antigen in a multivalent ACV containing PT antigen.

Using standard immunological analyses, we observed typical Th2 and Th1/17 responses in ACV and WCV immunized mice respectively. *B. pertussis* utilizes PT and ACT to facilitate survival in the host and most studies about the PT/ACT specific effects have been perform on cell cultures *in vitro*. Microarray analysis has been used to profile the lung transcriptome of mice challenged with *B. pertussis*^40,57^. Here, we sought to use RNAseq to investigate the overall gene expression profiles of the murine lung in response to *B. pertussis* challenge. Analysis of the lung transcriptome 1 day after *B. pertussis* challenged exhibited a distinct profile in mice vaccinated with WCV compared to ACV, RTX, or naïve infected mice. This response was consistent with our WCV cytokine profile demonstrating a strong pro-inflammatory response. Similarly, transcriptome data from others during early *B. pertussis* infection noted an increase in chemokines such as Cxcl2, Cxcl10, Cxcl3, Ccl3, Cxcl1, and Ccl4^57,58^. In Raeven *et al*, convalescent mice previously infected with *B. pertussis* exhibited higher expression of these chemokines compared to naïve mice, similar to our whole cell transcriptome where these chemokines are consistently higher than the naïve mice (Fig. 7 and Fig. 8). We observed the highest expression of these chemokines in the naïve and RTX mice groups, where these groups had a similar gene profile of genes belonging to the immune cell trafficking and cardiovascular disease annotations (Fig. 7). These findings, along with higher bacterial burden (Fig. 3) suggest that RTX-alum immunization does not provide sufficient protection.

Our overall transcriptomic profiling revealed thousands of significant gene expression changes (Fig. 6). After analyzing the neutrophil (Fig. 8), and innate gene changes (Supplementary Fig. 4), we then sought to specifically characterize B cell clone diversity. The RNAseq reads were re-processed with the MiXCR algorithm and we analyzed the abundance and diversity of VDJ clonotypes (Supplementary Fig 5). Besides the T cell response differences of the ACV and WCV (Th2 v. Th1/Th17) another significant difference between these two vaccines is the number of antigens. ACVs have 3-5 antigens (PT, FHA, PRN, FIM2/3) but the WCV hypothetically has ∼3,000 antigens. In light of this, it would be logical to hypothesize that the WCV would induce a vast antibody repertoire as measured by many VDJ clonotypes. However, we observed a greater diversity and abundance in the ACV group compared to the WCV at both early and late time-points (Supplementary Fig. S6). It is important to point out that this analysis was performed on total lung RNA. If we were to isolate B cells and sequence deeper we would expect to more thoroughly characterize the repertoire. These data are interesting but we do not know which clone types result in functional antibodies that protect against *B. pertussis*. Further analysis is required to bridge the gap between clonotypes and functional/protective antibodies.

Using NeCRE luc mice, we performed tracking neutrophil recruitment during a *B. pertussis* respiratory infection. These data demonstrate how analysis of cellular responses through *in vivo* imaging can be used to provide a quantitative parameter throughout a study. This model can be applied to other bacterial infection models or cancer tumor progression models where following the same mouse throughout a study would be beneficial. In this study, we also added next generation sequencing technology to expand upon the immunological findings. RNAseq analysis can be further employed to understand key cell populations and how they are impacted by both immunization and *B. pertussis* challenge. Our current goal is to continue to refine the murine challenge models with new technological approaches in order to facilitate formulation of new pertussis vaccines. The findings of this study suggest that by integrating RNAseq analysis with classic immunological techniques, it is possible to illuminate novel intricacies of vaccine induced immunity to *B. pertussis*.

## Methods

### Bacteria and culture conditions

*B. pertussis* strain UT25 (UT25Sm1) were cultured on Bordet-Gengou (BG) agar (1906) supplemented with 15% defibrinated sheep blood (Hemostat Laboratories) for 48 h at 36°C^59^. *B. pertussis* was then transferred from BG plates to three flasks of 12 ml of modified Stainer-Scholte liquid medium (SSM)^60^. SSM cultures were not supplemented with cyclodextrin (Heptakis(2,6-di-O-methyl)-β-cyclodextrin). SSM cultures were grown for ∼22 h at 36°C with shaking at 180 rpm until the OD600 reached 0.5 on a 1 cm path width spectrophotometer (Beckman Coulter DU 530). The cultures were then diluted to 1 × 10^9^/ml with SSM.

### Mouse Strains

All mouse strains used were bred in a specific pathogen-free experimental conditions within the Office of Laboratory Animal Resources vivarium at West Virginia University. Mice were aged 8 – 12 weeks, male and female sex mice were equally assigned to all vaccination groups. 129- Elane^tm1(cre)Roes^/H mice (Medical Research Council, London, UK) were crossed with FVB.129S6(B6)-Gt(ROSA)26Sortm1(luc)kael/J mice (Jackson labs; 0051225) resulting in NECre luc progeny. NECre luc mice were injected with CycLuc 1 lucferin (EMD Millipore, Darmstadt, Germany) and confirmed to be luminescent using Xenogen Lumina II^61^. Upon intraperitoneal injection of a luciferase substrate (luciferin), neutrophils emit luminescence that is detectable using a high sensitivity live animal imaging system (Xenogen IVIS Lumina II).

### Vaccines used in study and administration

All vaccines were formulated into 200 µl doses with the antigen content described below. 100 µl of INFANRIX (GSK) human vaccine (DTaP) which is 1/5 human dose of the vaccine, was mixed in 100 µl of PBS. The NIBSC WHO standard *Bordetella pertussis* whole-cell vaccine (NIBSC code 94/532) was received lyphophilzed and reconstituted in 1ml of PBS. At this concentration one human dose is 100 µl therefore, 20 µl was mixed with 180 µl of PBS, which is 1/5 of the human dose (66 µg of total protein). A truncated ACT protein (RTX) was purified as previously described ^22^ 5.6 ug of RTX was combined with 100 µl alum adjuvant (Alhydrogel®, InvivoGen) corresponding to 1 mg of aluminum hydroxide. All immunizations occurred by intraperitoneal injection. Unvaccinated mice received 200 µl of sterile PBS.

### Vaccination and Challenge with *Bordetella pertussis*

NECre luc mice were bred to ages ranging from (8-12 weeks). Mice were vaccinated at day 0, and then boosted 21 days later. Thirty-five days post initial vaccination, *B. pertussis* UT25 was grown as described above, and provided as a challenge dose at 2 × 10^7^ CFU in 20 µl. Mice were anesthetized by intraperitoneal injection of 200 µl of ketamine (6.7 mg/ml) and xylazine (1.3 mg/ml) in 0.9% saline. Two 10 µl doses of bacteria were administered through nasal inhalation into each nostril of the mouse. Mice from each of the groups were challenged WCV (8), ACV (8), RTX (6), and naïve control (PBS injection) (5).

### Collection of murine samples and determination of bacterial burden

On days 1,2,4,6, and 9 pc, mice were euthanized by intraperitoneal injection of pentobarbital and dissected in a biosafety cabinet under BSL-2 conditions. Blood was collected by cardiac puncture, serum was separated by centrifugation through a BD Microtainer SST blood collector (BD), or blood for complete blood cell counts were collected in BD Microtainer Tubes with K2EDTA (BD). Trachea and lungs were removed, placed in 1 ml PBS, and then homogenized. Trachea tissue was homogenized by a Brinkman Homogenizer (Polytron), while lung tissue was dissociated using a Dounce homogenizer (Kimble Chase). To determine viable *B. pertussis* in the nares, 1 ml of PBS was flushed through the nares and collected. In order to determine bacterial burden 100 µl of homogenate or nasal lavage was serial diluted in sterile PBS. Four 10 µl aliquots of each serial dilutions were plated on BG containing streptomycin (100 µg/ml) to ensure only UT25 *B. pertussis* grew on the plates. After 72 h at 36°C colony forming units (CFUs) were counted and the bacterial burden per tissue was calculated. Due to the serial dilutions plated our limit of detection was 10^3^ CFUs per ml or organ. CFUs from experimental groups at each time-point were compared to the naive (PBS injected) and challenged group by a two-tailed unpaired t test using the software package Prism 7 (GraphPad, La Jolla, CA). All experiments were performed in accordance with the National Institutes of Health Guide for the Care and Use of Laboratory Animals. All murine infection experiments were performed per protocols approved by the West Virginia University Institutional Animal Care and Use Committee (protocol number Damron 14- 1211).

### IVIS imaging

NECre luc mice received 83 mg/kg of CycLuc1 luciferin by IP injection (100 µl). Five minutes after injection mice were anesthetized with 3% isoflurane, mixed with oxygen from the XGI-8 gas anesthesia system supplied with a Xenogen IVIS Lumina II. Luminescent signals were acquired during a five-minute exposure. Acquisition was performed using Living Image 2.5 software (Xenogen). Images were acquired with a binning of 4. Following imaging, mice were either euthanized for dissection and extraction of tissue samples or returned to the vivarium. Luminescence was determined by quantification of photons emitted per second generated in each region of interest (ROI): whole mouse image or nasal cavity. The background of each image was subtracted from the ROI, background photons were defined by the average of two ROIs away from the mouse, for the same image. Data was expressed as relative fold change between target ROIs from experimental groups to an average of ROIs from control mice that were not vaccinated or challenged. Group comparisons were analyzed by one-way analysis of variance (ANOVA) followed by a Tukey’s multiple-comparison test using Prism 7.

### Preparation of tissue and flow cytometry analysis

Blood, lung, and cells isolated from nasal lavage were analyzed by flow cytometry. A 100 µl sample was removed from lung homogenate of all samples and filtered through a 70 µm cell strainer, then centrifuged at 1000 × *g* for 5 mins to pellet cells. Supernatant was removed, and RBC lysis buffer (BD Pharm lysis) added incubated at 37°C for 2 min, then pelleted using the same centrifugation conditions. Cells were resuspended in 500 µl of PBS + 1% FBS, 100µl was aliquoted for antibody staining. After cardiac puncture blood was placed in EDTA containing tube (BD Bioscience), RBC were lysed using Pharmlyse (BD Biosciences) with a 15 min room temperature incubation and then prepared in a similar manner to other tissues. All cell suspension samples were incubated on ice in PBS and 1% FBS for blocking. Antibodies against specific cell surface markers: PE-conjugated GR-1 (BD, 553128) Alexa Fluor 700-conjugated CD11b (Biolegend, 101222) were added to cell suspensions and incubated in the dark for 1 h at 4ᵒC. Lung, blood, and nasal wash suspensions were pelleted, and resuspended in PBS prior to analysis. Samples were read using LSR Fortessa (BD), and analyzed using FlowJo v10 (FlowJo, LLC). PMNs were classified as CD11b^+^Gr-1^+^ single, live cells.

### Cytokine Quantification

Lung homogenates were pelleted by centrifugation and then supernatant was collected and stored at −80°C until analysis. Concentration of cytokines in the lungs of vaccinated and challenged mice were determined by quantitative sandwich immunoassays, Meso Scale Discovery (Rockville, MD) V-PLEX Proinflammatory Panel (K15048G-1) and Mouse IL-17 Ultra-Sensitive kits (K152ATC-1), following manufacturer’s instructions. Data was analyzed by one-way ANOVA, with a Tukey’s multiple-comparison test for each time-point.

### Serology

Vaccinated and challenged mouse serological responses to RTX, and PT were determined by qualitative ELISA. High-binding 96-well ELISA plates were coated overnight at 4°C with 50 µl of purified RTX or Pt in PBS at a concentration of 1 µg/mL. Purified *B. pertussis* antigens, were obtained from Dr. Jennifer Maynard. Serum samples from NVNC, PBS, ACV, and WCV serum titers were analyzed for PT, as the RTX group had no PT in vaccine. Samples from NVNC, PBS, WCV, and RTX were analyzed for serum antibody titers to RTX, as ACV would not be expected to have RTX titers because no RTX was included in vaccine. Plates were then washed with PBS + 1% Tween 20 (PBS-T), then blocked with 5% milk in PBS-T for 1 hour at room temperature. Sera were diluted to a concentration in the linear dose range for each antigen. Plates were incubated for 2 h at 37⁰C with serial diluted serum samples from vaccinated groups. Following 3 PBS-T washes, 1:4000 goat anti-mouse IgG-AP (Southern Biotech), secondary antibody was added and incubated 1 h at 37⁰C. Plates were washed, then developed for 30 min with 100 µl p-nitrophenyl phosphate. Colorimetric signal was measured using Spectramax i3 (Molecular Devices) at 450nm. An average of blanks was subtracted from all absorbances and used as a baseline detection limit. The minimum detection limit above the baseline was analyzed by one-way ANOVA, with a Tukey’s multiple-comparison test for each time-point using Prism 7.

### Isolation of Lung RNA, illumina library preparation, and sequencing

Lung RNA was prepared similar to previously reported^62^ with the following modifications. The freshly isolated NECre luc mice lungs were homogenized and RNA was prepared immediately using RNeasy purification kit (Qiagen). Each lung was placed in 1 ml of sterile PBS and then 2 ml of TE lysozyme (1mg/ml) was added and allowed to incubate for 10 min on ice. 2 ml of RLT buffer was added and incubated for another 10 min on ice. The homogenate was then pushed through a syringe needle. The homogenates were pelleted by centrifugation at 20,800g (max speed microfuge) for 10 min. The supernatant was extracted and then 2.8ml of 100% EtOH was added to each tube. This supernatant of each mouse sample was then disturbed to four RNeasy tubes for RNA isolation. The RNA was eluted and pooled into one sample per mouse. The resulting RNA was quantified on a Qubit 3.0 (ThermoFisher) with the high intensity assay kit. Next, the RNA integrity was assessed using Agilent BioAnalyzer RNA Pico chip. All samples were then submitted Ribo-zero rRNA depletion (illumina) and reassessed for RNA integrity. rRNA depleted mRNA samples were then fragmented and prepared into libraries using illumina ScriptSeq Complete Gold (Epidemiology). Libraries were checked for quality control with KAPA qPCR QC assay (KAPA Biosystems). The libraries (33 total) were then sequenced on an illumina HiSeq at the Marshall University Genomics Core facility on 2 lanes of 2×50*B. pertussis*. Sequencing data were deposited to the Sequence Read Archive (SRA) and are available under the reference number SRA587785, BioProject number PPJNA394758.

### RNAseq bioinformatics analyses

The reads were analyzed using the software CLC Genomics workbench 9.5. *Mus musculus* genome was downloaded from NCBI (version GRCm38.78). Reads were mapped against the genome using the following settings for mapping: mismatch cost = 2, insertion cost = 3, deletion cost = 3, length fraction = 0.8, similarity fraction = 0.8. RPKM values were generated using default parameters for CLC Genomics. On average ∼20 million reads were obtained for each sample and with stringent parameters ∼73% mapped to the murine genome with our mapping parameters. Fold changes in gene expression and statistical analyses were performed using an Extraction of Differential Gene Expression (EDGE) test p value. Expression data for each gene was considered significant if the p value was less than 0.05. Venn diagrams were generated using Venny 2.1^63^. Fold-change of gene expression data was plotted relative to non-challenged control groups at each time-point. Supplementary Table 8 contains the exported gene expression analysis worksheets and statistical analyses. Gene list were created using GO terms acquired though AmiGO 2 database including neutrophil (CL:000075), neutrophil activation (GO:0042119), neutrophil migration (GO:1990266), innate immune response (GO: 0045087), T-helper 1 type immune response (GO:0042088), type 2 immune response (GO:0042092), and T-helper 17 type immune response (GO:0072538)^64^.

### Pathway Enrichment Analysis

Ingenuity Pathway Analysis (Qiagen) was utilized to map lung transcriptomes to biological and disease functions. Lung transcriptomes were loaded to IPA, and comparative analyses were performed at early and late time-points. Gene expression fold-changes were mapped to higher-order disease and function defined by the IPA knowledge base, and a gene enrichment analysis was performed on early and late time-points. A Fisher Exact T-test was used to determine statistically significant (p < 0.05) p-values of the functions that made up the higher-order disease category and represented as a range of p-values. The relative fold-change of these identified genes was determined, and the 10 highest fold-changes were represented as heat maps.

### Immunoglobulin and T-cell receptor profiling

B cell clones were identified using MiXCR software (MiLabratory), capable of generating quantitated clonotypes of immunoglobulins^48^. The same paired-end Illumina sequenced reads mentioned above were merged using concatenation, then imported into MiXCR software. Merged reads were aligned to each other to generate clonotypes of based on VDJ segment regions of unique immunoglobins specific to each sample. Clone data for each sample was grouped based on vaccine received, and time-point. Clonotypes from each sample were then separated into T and B cells, based on T-cell receptor or B-cell receptor specific sequences. Prepared data files were then imported into VDJtools (MiLabratory), for data representation according to established protocol^65^. Briefly, data was represented to show V-J diversity, and quantify unique clonotypes using dendrograms and chord diagrams.

## Supporting information

Boehm et al Supplemental Information

## Acknowledgements

*B. pertussis* strain UT25 was kindly provided by Dr. Sandra Armstrong (University of Minnesota). The NeCre luc mice were developed by Ian Glomski and we thank him for originally providing the mice and developing the imaging methodologies. The Elane mice were kindly provided by MRC Harwell (Oxfordshire, UK). RTX antigen was graciously purified by Andrea DiVenere, University of Texas at Austin. We would like to thank the following WVU facilities: Genomics Core, Office of Laboratory Animal Resources (OLAR) for support with the murine studies, Animal Models and Imaging (U54 GM104942) for support IVIS imaging, the flow cytometry and single cell (S10 OD016165). We would like to acknowledge Kathy Brundage for flow cytometry support, Sarah McLaughlin for IVIS support and Ryan Percifield for next generation sequencing library preparation. D.T.B. was supported by the WVU HSC Office of Research and Graduate Education, graduate student fellowship from the West Virginia NASA Space Grant Consortium, and the Jennifer Gossling Fellowship. Mackenna Boone was supported by a WV-InBRE summer fellowship. Additional support was provided by NIH/NIAID grant RO1 AI1018000 (E.L.H.) and NIH/NIAID grant RO1 A1122753 (J.A.M). This work was also supported by funding from National Institutes of Health HHSN272201200005C-416476 and laboratory startup funds from West Virginia University to F.H.D. The Marshall University CORE facilities and RNA sequencing were funded by the WV InBRE grant P20 GM103434.

## Author contributions

All authors participated in the composition and review of the manuscript. D.T.B. designed experiments, performed IVIS imaging, raised and dissected mice, analyzed immunological data, mapped NGS read data and calculated expression analysis. M.E.V. developed flow cytometry panels, and performed analyses. T.W. coordinated murine trials, and prepared RNA for NGS analysis. E.S.N. and E.S.K. analyzed RNAseq and developed visualizations. J.M.H. performed serological analysis. C.E., M.B., S.B., K.D., J.B., and M.E prepared and analyzed samples to measure correlates of protection on each experimental day. J.M. provided the RTX antigen. J.M., E.L.H, M.B. and F.H.D. designed the overall strategy of the studies. M.B. and F.H.D. formulated vaccines, directed experiment days, performed dissections, and analyzed RNAseq data.

## Supplemental Information

Supplemental information is available under Boehm et al Supplemental Information.

## References

1 Paddock CD, Sanden GN, Cherry JD, Gal AA, Langston C, Tatti KM, et al. Pathology and Pathogenesis of Fatal *Bordetella pertussis* Infection in Infants. Clin Infect Dis 2008;47:328–38. https://doi.org/10.1086/589753.

2 Andreasen C, Powell DA, Carbonetti NH. Pertussis toxin stimulates IL-17 production in response to Bordetella pertussis infection in mice. PLoS One 2009;4:e7079. https://doi.org/10.1371/journal.pone.0007079.

3 Carbonetti NH. Pertussis toxin and adenylate cyclase toxin: Key virulence factors of Bordetella pertussis and cell biology tools. Future Microbiol 2010;5:455–69. https://doi.org/10.2217/fmb.09.133.

4 Carbonetti NH. Contribution of pertussis toxin to the pathogenesis of pertussis disease. Pathog Dis 2015;73:ftv073. https://doi.org/10.1093/femspd/ftv073.

5 Carbonetti NH, Artamonova G V., Andreasen C, Bushar N. Pertussis toxin and adenylate cyclase toxin provide a one-two punch for establishment of Bordetella pertussis infection of the respiratory tract. Infect Immun 2005;73:2698–703. https://doi.org/10.1128/IAI.73.5.2698-2703.2005.

6 Sebo P, Osicka R, Masin J. Adenylate cyclase toxin-hemolysin relevance for pertussis vaccines. Expert Rev Vaccines 2014;13:1215–27. https://doi.org/10.1586/14760584.2014.944900.

7 Mattoo S, Cherry JD. Molecular pathogenesis, epidemiology, and clinical manifestations of respiratory infections due to Bordetella pertussis and other Bordetella subspecies. Clin Microbiol Rev 2005;18:326–82. https://doi.org/10.1128/CMR.18.2.326-382.2005.

8 Gold MS. Hypotonic-Hyporesponsive Episodes Following Pertussis Vaccination. Drug Saf 2002;25:85–90. https://doi.org/10.2165/00002018-200225020-00003.

9 Donnelly S, Loscher CE, Lynch MA, Mills KHG. Whole-cell but not acellular pertussis vaccines induce convulsive activity in mice: Evidence of a role for toxin-induced interleukin-1?? in a new murine model for analysis of neuronal side effects of vaccination. Infect Immun 2001;69:4217–23. https://doi.org/10.1128/IAI.69.7.4217-4223.2001.

10 Ross PJ, Sutton CE, Higgins S, Allen AC, Walsh K, Misiak A, et al. Relative contribution of Th1 and Th17 cells in adaptive immunity to Bordetella pertussis: towards the rational design of an improved acellular pertussis vaccine. PLoS Pathog 2013;9:e1003264. https://doi.org/10.1371/journal.ppat.1003264.

11 Mills KHG, Ryan M, Ryan E, Mahon BP. A murine model in which protection correlates with pertussis vaccine efficacy in children reveals complementary roles for humoral and cell-mediated immunity in protection against Bordetella pertussis. Infect Immun 1998;66:594–602. https://doi.org/10.1371/journal.ppat.1003264.

12 Warfel JM, Merkel TJ. Bordetella pertussis infection induces a mucosal IL-17 response and long-lived Th17 and Th1 immune memory cells in nonhuman primates. Mucosal Immunol 2013;6:787–96. https://doi.org/10.1038/mi.2012.117.

13 Carbonetti NH. Pertussis leukocytosis: mechanisms, clinical relevance and treatment. Pathog Dis 2016;74:ftw087. https://doi.org/10.1093/femspd/ftw087.

14 Fedele G, Spensieri F, Palazzo R, Nasso M, Cheung GYC, Coote JG, et al. Bordetella pertussis commits human dendritic cells to promote a Th1/Th17 response through the activity of adenylate cyclase toxin and MAPK-pathways. PLoS One 2010;5:e8734. https://doi.org/10.1371/journal.pone.0008734.

15 Mills KHG, Ross PJ, Allen AC, Wilk MM. Do we need a new vaccine to control the re-emergence of pertussis? Trends Microbiol 2014;22:49–52. https://doi.org/10.1016/j.tim.2013.11.007.

16 Klein NP, Bartlett J, Fireman B, Baxter R. Waning Tdap Effectiveness in Adolescents. Pediatrics 2016;137:e20153326–e20153326. https://doi.org/10.1542/peds.2015-3326.

17 Warfel JM, Beren J, Merkel TJ. Airborne transmission of bordetella pertussis. J Infect Dis 2012;206:902–6. https://doi.org/10.1093/infdis/jis443.

18 Althouse BM, Scarpino S V. Asymptomatic transmission and the resurgence of *Bordetella pertussis*. BMC Med 2015;13:146. https://doi.org/10.1186/s12916-015-0382-8.

19 Betsou F, Sebo P, Guiso N. CyaC-mediated activation is important not only for toxic but also for protective activities of Bordetella pertussis adenylate cyclase-hemolysin. Infect Immun 1993;61:3583–9.

20 Cheung GYC, Xing D, Prior S, Corbel MJ, Parton R, Coote JG. Effect of different forms of adenylate cyclase toxin of Bordetella pertussis on protection afforded by an acellular pertussis vaccine in a murine model. Infect Immun 2006;74:6797–805. https://doi.org/10.1128/IAI.01104-06.

21 Guiso N, Szatanik M, Rocancourt M. Protective activity of Bordetella adenylate cyclase-hemolysin against bacterial colonization. Microb Pathog 1991;11:423–31. https://doi.org/10.1016/0882-4010(91)90038-C.

22 Wang X, Maynard JA. The Bordetella adenylate cyclase repeat-in-toxin (RTX) domain is immunodominant and elicits neutralizing antibodies. J Biol Chem 2015;290:3576–91. https://doi.org/10.1074/jbc.M114.585281.

23 Harvill ET, Cotter PA, Miller JF. Pregenomic comparative analysis between Bordetella bronchiseptica RB50 and Bordetella pertussis Tohama I in murine models of respiratory tract infection. Infect Immun 1999;67:6109–18.

24 Kirimanjeswara GS, Agosto LM, Kennett MJ, Bjornstad ON, Harvill ET. Pertussis toxin inhibits neutrophil recruitment to delay antibody-mediated clearance of Bordetella pertussis. J Clin Invest 2005;115:3594–601. https://doi.org/10.1172/JCI24609.

25 Andreasen C, Carbonetti NH. Role of neutrophils in response to bordetella pertussis infection in mice. Infect Immun 2009;77:1182–8. https://doi.org/10.1128/IAI.01150-08.

26 Weiss AA, Mary GMS. Lethal infection by Bordetella pertussis mutants in the infant mouse model. Infect Immun 1989;57:3757–64.

27 Goodwin MS, Weiss AA. Adenylate cyclase toxin is critical for colonization and pertussis toxin is critical for lethal infection by Bordetella pertussis in infant mice. Infect Immun 1990;58:3445–7.

28 Locht C, Coutte L, Mielcarek N. The ins and outs of pertussis toxin n.d. https://doi.org/10.1111/j.1742-4658.2011.08237.x.

29 Froehlich. Beitrag zur pathologie des Keuchhustens. Jahrb Fur Kinderheilkd 1897;44:.

30 Meunier H. De la leucocytose dans la coqueluche. CR Soc Biol 1898;50:.

31 Morse SI. STUDIES ON THE LYMPHOCYTOSIS INDUCED IN MICE BY BORDETELLA PERTUSSIS. J Exp Med 1965;121:49–68. https://doi.org/10.1084/jem.121.1.49.

32 Morse SI, Riester SK. Studies on the leukocytosis and lymphocytosis induced by Bordetella pertussis. I. Radioautographic analysis of the circulating cells in mice undergoing pertussis-induced hyperleukocytosis. J Exp Med 1967;125:401–8. https://doi.org/10.1084/jem.125.3.401.

33 Weiner ZP, Ernst SM, Boyer AE, Gallegos-Candela M, Barr JR, Glomski IJ. Circulating lethal toxin decreases the ability of neutrophils to respond to Bacillus anthracis. Cell Microbiol 2014;16:504–18. https://doi.org/10.1111/cmi.12232.

34 Higgs R, Higgins SC, Ross PJ, Mills KHG. Immunity to the respiratory pathogen Bordetella pertussis. Mucosal Immunol 2012;5:485–500. https://doi.org/10.1038/mi.2012.54.

35 Ross PJ, Sutton CE, Higgins S, Allen AC, Walsh K, Misiak A, et al. Relative contribution of Th1 and Th17 cells in adaptive immunity to *Bordetella pertussis* : towards the rational design of an improved acellular pertussis vaccine. PLoS Pathog 2013;9:e1003264. https://doi.org/10.1371/journal.ppat.1003264.

36 Alvarez Hayes J, Erben E, Lamberti Y, Principi G, Maschi F, Ayala M, et al. Bordetella pertussis iron regulated proteins as potential vaccine components. Vaccine 2013;31:3543–8. https://doi.org/10.1016/j.vaccine.2013.05.072.

37 Sawal M, Cohen M, Irazuzta JE, Kumar R, Kirton C, Brundler M-A, et al. Fulminant pertussis: A multi-center study with new insights into the clinico-pathological mechanisms. Pediatr Pulmonol 2009;44:970–80. https://doi.org/10.1002/ppul.21082.

38 Pierce C, Klein N, Peters M. Is leukocytosis a predictor of mortality in severe pertussis infection? Intensive Care Med 2000;26:1512–4.

39 Alves-Filho JC, Freitas A, Souto FO, Spiller F, Paula-Neto H, Silva JS, et al. Regulation of chemokine receptor by Toll-like receptor 2 is critical to neutrophil migration and resistance to polymicrobial sepsis. Proc Natl Acad Sci U S A 2009;106:4018–23. https://doi.org/10.1073/pnas.0900196106.

40 Raeven RHM, Van Der Maas L, Tilstra W, Uittenbogaard JP, Bindels THE, Kuipers B, et al. Immunoproteomic Profiling of Bordetella pertussis Outer Membrane Vesicle Vaccine Reveals Broad and Balanced Humoral Immunogenicity. J Proteome Res 2015;14:2929–42. https://doi.org/10.1021/acs.jproteome.5b00258.

41 Starost LJ, Karassek S, Sano Y, Kanda T, Kim KS, Dobrindt U, et al. Pertussis Toxin Exploits Host Cell Signaling Pathways Induced by Meningitis-Causing E. coli K1-RS218 and Enhances Adherence of Monocytic THP-1 Cells to Human Cerebral Endothelial Cells. Toxins (Basel) 2016;8:291. https://doi.org/10.3390/toxins8100291.

42 Cotoi OS, Dunér P, Ko N, Hedblad B, Nilsson J, Björkbacka H, et al. Plasma S100A8/A9 correlates with blood neutrophil counts, traditional risk factors, and cardiovascular disease in middle-aged healthy individuals. Arterioscler Thromb Vasc Biol 2014;34:202–10. https://doi.org/10.1161/ATVBAHA.113.302432.

43 Vogl T, Ludwig S, Goebeler M, Strey A, Thorey IS, Reichelt R, et al. MRP8 and MRP14 control microtubule reorganization during transendothelial migration of phagocytes. Blood 2004;104:4260–8. https://doi.org/10.1182/blood-2004-02-0446.

44 Kumar H, Kumagai Y, Tsuchida T, Koenig PA, Satoh T, Guo Z, et al. Involvement of the NLRP3 Inflammasome in Innate and Humoral Adaptive Immune Responses to Fungal-Glucan. J Immunol 2009;183:8061–7. https://doi.org/10.4049/jimmunol.0902477.

45 Gabrilovich DI. The Neutrophils : New Outlook for Old Cells. Singapore, UNITED STATES: Imperial College Press; 2014.

46 Mahon BP, Sheahan BJ, Griffin F, Murphy G, Mills KH. Atypical disease after Bordetella pertussis respiratory infection of mice with targeted disruptions of interferon-gamma receptor or immunoglobulin mu chain genes. J Exp Med 1997;186:1843–51.

47 Kirimanjeswara GS, Mann PB, Harvill ET. Role of antibodies in immunity to Bordetella infections. Infect Immun 2003;71:1719–24. https://doi.org/10.1128/iai.71.4.1719-1724.2003.

48 Bolotin DA, Poslavsky S, Mitrophanov I, Shugay M, Mamedov IZ, Putintseva E V, et al. MiXCR: software for comprehensive adaptive immunity profiling. Nat Methods 2015;12:380–1. https://doi.org/10.1038/nmeth.3364.

49 Xu JL, Davis MM. Diversity in the CDR3 region of V(H) is sufficient for most antibody specificities. Immunity 2000;13:37–45. https://doi.org/10.1016/S1074-7613(00)00006-6.

50 Nguyen AW, Wagner EK, Laber JR, Goodfield LL, Smallridge WE, Harvill ET, et al. A cocktail of humanized anti-pertussis toxin antibodies limits disease in murine and baboon models of whooping cough. Sci Transl Med 2015;7:316ra195. https://doi.org/10.1126/scitranslmed.aad0966.

51 Im S-Y, Wiedmeier SE, Cho B-H, Lee DG, Beigi M, Daynes RA. Dual effects of pertussis toxin on murine neutrophils in vivo. Inflammation 1989;13:707–26. https://doi.org/10.1007/BF00914314.

52 Winter K, Zipprich J, Harriman K, Murray EL, Gornbein J, Hammer SJ, et al. Risk Factors Associated With Infant Deaths From Pertussis: A Case-Control Study. Clin Infect Dis 2015;61:1099–106. https://doi.org/10.1093/cid/civ472.

53 Heininger U, Klich K, Stehr K, Cherry JD. Clinical findings in Bordetella pertussis infections: results of a prospective multicenter surveillance study. Pediatrics 1997;100:E10. https://doi.org/10.1542/peds.100.6.e10.

54 Guiso N, Rocancourt M, Szatanik M, Alonso JM. Bordetella adenylate cyclase is a virulence associated factor and an immunoprotective antigen. Microb Pathog 1989;7:373–80. https://doi.org/10.1016/0882-4010(89)90040-5.

55 Torres CA, Iwasaki A, Barber BH, Robinson HL. Differential dependence on target site tissue for gene gun and intramuscular DNA immunizations. J Immunol 1997;158:4529–32.

56 Bassinet L, Fitting C, Housset B, Cavaillon JM, Guiso N. Bordetella pertussis adenylate cyclase-hemolysin induces interleukin-6 secretion by human tracheal epithelial cells. Infect Immun 2004;72:5530–3. https://doi.org/10.1128/IAI.72.9.5530-5533.2004.

57 Raeven RHM, Brummelman J, Pennings JLA, Nijst OEM, Kuipers B, Blok LER, et al. Molecular signatures of the evolving immune response in mice following a Bordetella pertussis infection. PLoS One 2014;9:e104548. https://doi.org/10.1371/journal.pone.0104548.

58 Moreno G, Errea A, Van Maele L, Roberts R, Léger H, Sirard JC, et al. Toll-like receptor 4 orchestrates neutrophil recruitment into airways during the first hours of Bordetella pertussis infection. Microbes Infect 2013;15:708–18. https://doi.org/10.1016/j.micinf.2013.06.010.

59 Brickman TJ, Armstrong SK. The ornithine decarboxylase gene odc is required for alcaligin siderophore biosynthesis in *Bordetella* spp.: Putrescine is a precursor of alcaligin. J Bacteriol 1996;178:54–60. https://doi.org/10.1128/jb.178.1.54-60.1996.

60 Stainer DW, Scholte MJ. A Simple Chemically Defined Medium for the Production of Phase I *Bordetella pertussis*. J Gen Microbiol 1970;63:211–20. https://doi.org/10.1099/00221287-63-2-211.

61 Evans MS, Chaurette JP, Adams ST, Reddy GR, Paley MA, Aronin N, et al. A synthetic luciferin improves bioluminescence imaging in live mice. Nat Methods 2014;11:393–5. https://doi.org/10.1038/nmeth.2839.

62 Damron FH, Oglesby-Sherrouse AG, Wilks A, Barbier M. Dual-seq transcriptomics reveals the battle for iron during *Pseudomonas aeruginosa* acute murine pneumonia. Sci Rep 2016;6:. https://doi.org/Artn 39172 10.1038/Srep39172.

63 Olivers JC. Venny. An interactive tool for comparing lists with Venn’s diagrams n.d.

64 Carbon S, Ireland A, Mungall CJ, Shu S, Marshall B, Lewis S, et al. AmiGO: online access to ontology and annotation data. Bioinformatics 2009;25:288–9. https://doi.org/10.1093/bioinformatics/btn615.

65 Shugay M, Bagaev D V, Turchaninova MA, Bolotin DA, Britanova O V, Putintseva E V, et al. VDJtools: Unifying Post-analysis of T Cell Receptor Repertoires. PLoS Comput Biol 2015;11:e1004503. https://doi.org/10.1371/journal.pcbi.1004503.

