## Supplementary material for "Characterizing the innate and adaptive responses of immunized mice to *Bordetella pertussis* infection using *in vivo* imaging and transcriptomic analysis": Boehm et al Supplemental Information

Short title: neutrophil responses to pertussis in mice

Dylan T. Boehm[a], Melinda E. Varney[a], Ting Y. Wong[a], Evan S. Nowak[a], Emel Sen-Kilic[a], Jesse Hall[a], Shelby D. Bradford[a], Katherine DeRoos[a], Justin Bevere[a], Matthew Epperly[a], Jennifer A. Maynard[b], Erik L. Hewlett[c], Mariette Barbier[a], and F. Heath Damron[a*]

[a] Department of Microbiology, Immunology, and Cell Biology, West Virginia University, Morgantown, WV, USA

[b] Department of Chemical Engineering, University of Texas at Austin, Austin, TX 78712, USA

[c] Department of Medicine, Division of Infectious Diseases and International Health, University of Virginia, Charlottesville, Virginia, USA

*Corresponding author

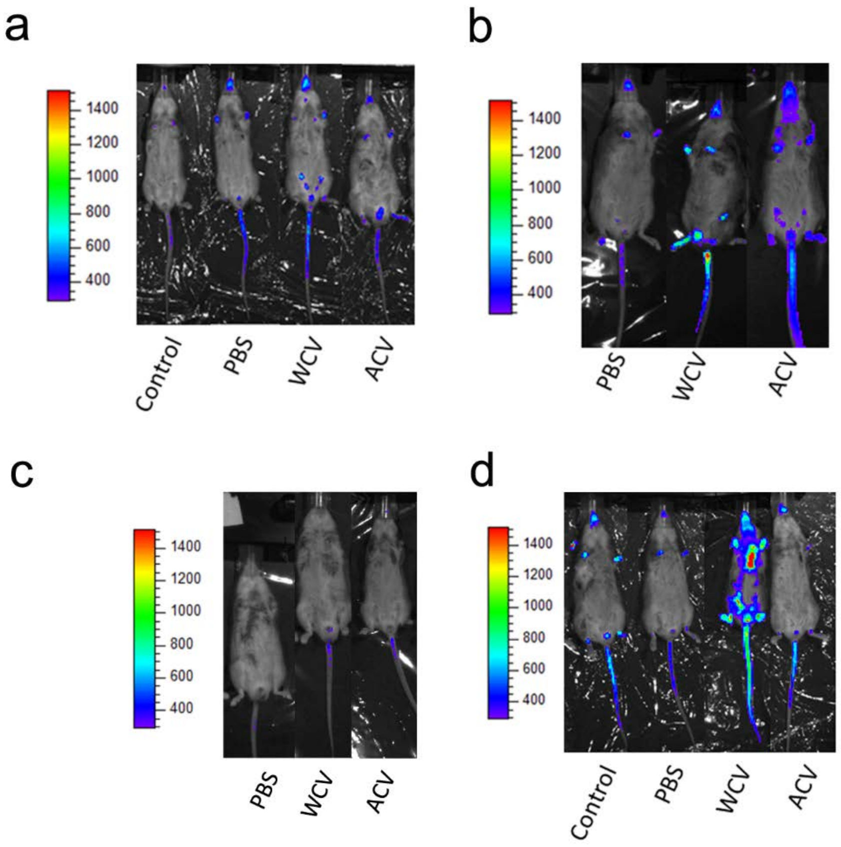


**Supplementary Fig. 1: IVIS imaging of naïve and immunized NECre luc mice post challenge with *Bp* as image was acquired from Xenogen IVIS Lumina II.** Images at day (a) 1, (b) 2, (c) 4, and (d) 6 – WCV or 9 – Control (Non-vaccinated/ no challenge), PBS, and ACV. Relative luminescence was quantified based on region of interest luminescence in these non-adjusted images.

**Supplementary Table 1. Compositions of vaccines of this study**

| | Vaccine groups | | |
|---|---|---|---|
| **Vaccine component*** | **aP (1/5th human dose)** | **wP**** | **RTX + alum** |
| Pertussis Toxoid*** | 5 | 0.4 | 0 |
| Filamentous Hemagglutinin | 5 | 3.5 | 0 |
| Pertactin | 1.6 | 0.3 | 0 |
| RTX | 0 | 0.2 | 10 |
| Aluminum hydroxide | 125 | 0 | 1 (mg) |
| Other antigens/adjuvants | 0 | 62 | 0 |

*All masses of antigens or adjuvant are indicated in µg

** This estimate is based on the number of peptides identified for each antigen in the wP. The percentage is then used to estimate the potential mass based on the fact that the wP dose used in this study contained 66 µg of total protein.

*** For total pertussis toxin, the sum of PtxA,B,C,D peptides was combined.

**Supplementary Table 2. Statistical analysis from Figure 3abc.**

| | Day 1 | | Day 2 | | Day 4 | |
|---|---|---|---|---|---|---|
| Trachrea | ACV | 0.0524 | WCV | 0.0341 | | |
| | WCV | 0.059 | | | | |
| | RTX | 0.0729 | | | | |
| Nasal Lavage | ACV | 0.0984 | ACV | 0.0764 | ACV | 0.0567 |
| | | | WCV | 0.0209 | WCV | 0.0567 |

Unpaired two-tailed T tests to PBS challenged group

**Supplementary Table 3: Relevant statistical differences in relative fold changes of neutrophil luminescence of *Bp* challenged mice.** Luminescence was analyzed by one-way ANOVA with a Tukey's multiple comparison test at each time point.

| | Day 1 | P value | Day 2 | P value | Day 3 | P value | Day 4 | P value | Day 6 | P value | Day 9 | P value |
|---|---|---|---|---|---|---|---|---|---|---|---|---|
| Whole Mouse | non-infected vs. PBS | 0.0002 | non-infected vs. PBS | 0.0004 | non-infected vs. PBS | 0.002 | non-infected vs. ACV | 0.013 | non-infected vs. WCV | 0.0001 | non-infected vs. PBS | 0.0006 |
| | non-infected vs. WCV | <0.0001 | non-infected vs. WCV | 0.0002 | non-infected vs. WCV | <0.0001 | | | non-infected vs. ACV | 0.0035 | non-infected vs. RTX | 0.0076 |
| | non-infected vs. ACV | 0.0004 | non-infected vs. ACV | 0.0003 | non-infected vs. ACV | 0.0003 | | | non-infected vs. RTX | 0.0042 | PBS vs. ACV | <0.0001 |
| | non-infected vs. RTX | 0.0007 | PBS vs. RTX | 0.0009 | non-infected vs. RTX | 0.0001 | | | | | ACV vs. RTX | 0.0003 |
| | PBS vs. WCV | 0.0007 | WCV vs. RTX | 0.0004 | PBS vs. WCV | 0.01 | | | | | | |
| | WCV vs. ACV | 0.0004 | ACV vs. RTX | 0.0006 | WCV vs. ACV | 0.0102 | | | | | | |
| | WCV vs. RTX | 0.0004 | | | | | | | | | | |
| Nasal Cacity | Day 1 | P value | Day 2 | P value | Day 3 | P value | Day 4 | P value | Day 6 | P value | Day 9 | P value |
| | non-infected vs. PBS | 0.0002 | non-infected vs. PBS | <0.0001 | non-infected vs. PBS | 0.0017 | non-infected vs. ACV | 0.0047 | non-infected vs. PBS | 0.0247 | non-infected vs. PBS | 0.0132 |
| | non-infected vs. WCV | <0.0001 | non-infected vs. WCV | <0.0001 | non-infected vs. WCV | <0.0001 | | | non-infected vs. WCV | 0.0004 | | |
| | non-infected vs. ACV | 0.0006 | non-infected vs. ACV | 0.0005 | non-infected vs. ACV | 0.0004 | | | non-infected vs. ACV | 0.0009 | | |
| | non-infected vs. RTX | 0.0005 | non-infected vs. RTX | 0.0005 | non-infected vs. RTX | 0.0002 | | | non-infected vs. RTX | 0.0038 | | |
| | WCV vs. ACV | 0.0262 | | | WCV vs. ACV | 0.0314 | | | | | | |

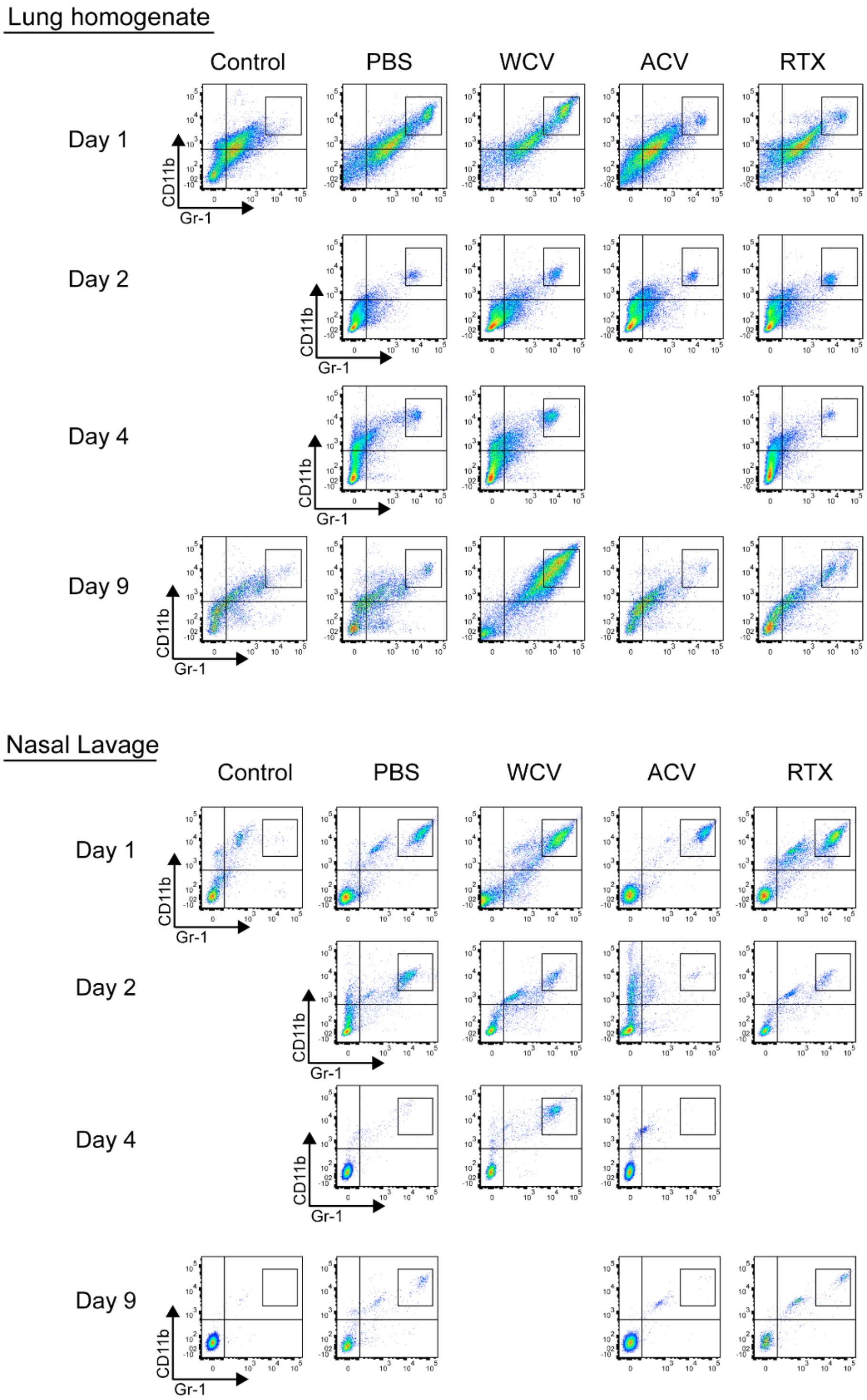


**Supplementary Fig. 2: Representative images and gating strategy for neutrophils from lung and nasal lavage tissues.** Neutrophils were gated as live, single cells (data not shown), quadrants demonstrate positively labeled cells determined by single marker flow cytometry controls. Region of interest shows neutrophil population ($Gr\text{-}1^{+}CD11b^{+}$). (a) Lung and (b) nasal lavage representatives.

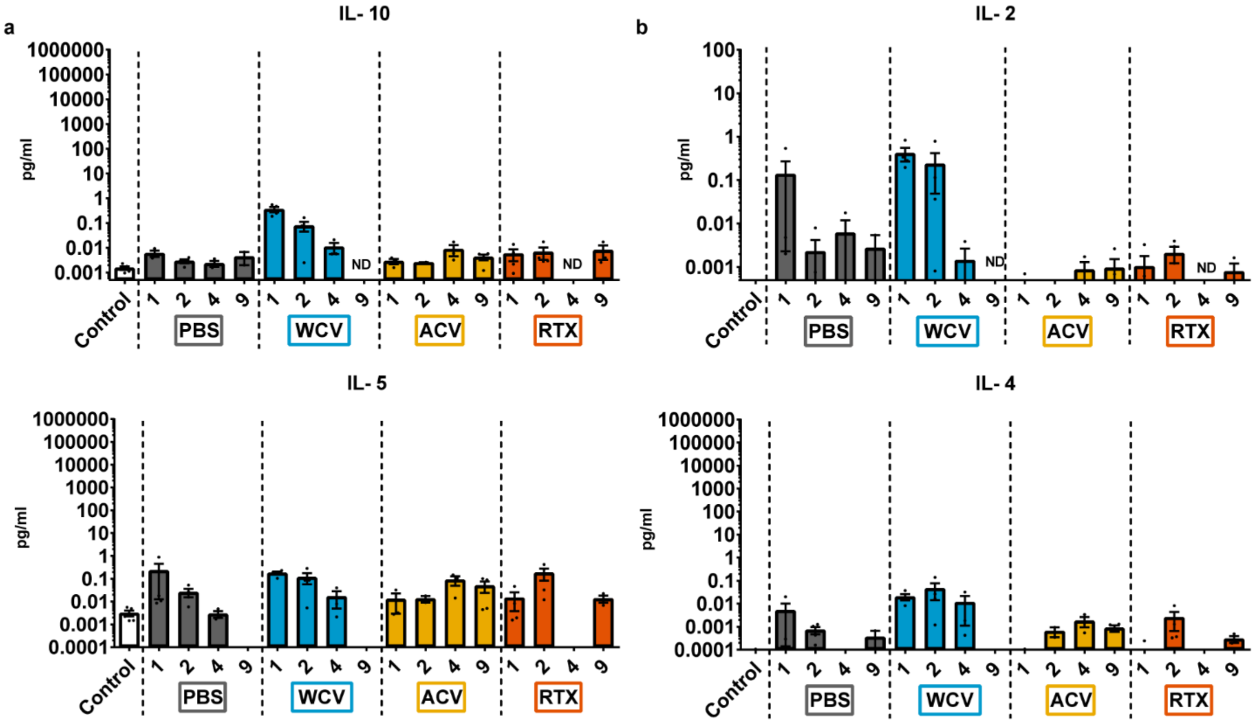


**Supplementary Fig. 3: Analysis of Treg and Th2 associated cytokines from lungs of naïve and immunized NECre luc mice post challenge with *Bp*.** Cytokines were analyzed at days 1, 2, 4 and 9 pc, then quantified using electrochemiluminescence immunoassays. (a) Cytokines IL-10 and IL-5 associated with Treg immune response. (b) Cytokines IL-2 and IL-4 associated with a Th2 immune response. ND: Sample not determined.

**Supplementary Table 4: Relevant statistical differences between cytokine production in lungs of immunized and *Bp* challenged NECre luc mice**. Samples were analyzed by one-way ANOVA with a Tukey's multiple comparison test at each time point.

| | Day 1 | | Day 2 | |
|---|---|---|---|---|
| | | P value | | P value |
| IL-6 | NV/NC vs. WCV | 0.0251 | | |
| | WCV vs. RTX | 0.0361 | | |
| IFN-γ | NV/NC vs. WCV | 0.0175 | NV/NC vs. WCV | 0.0022 |
| | PBS vs. WCV | 0.0261 | PBS vs. WCV | 0.004 |
| | WCV vs. RTX | 0.0408 | WCV vs. ACV | 0.0162 |
| | WCV vs. ACV | 0.0257 | WCV vs. RTX | 0.0039 |
| TNF-α | NV/NC vs. WCV | 0.0232 | | |
| | WCV vs. RTX | 0.033 | | |
| IL-1β | | | NV/NC vs. WCV | 0.0044 |
| | | | PBS vs. WCV | 0.0072 |
| | | | WCV vs. ACV | 0.0288 |
| | | | WCV vs. RTX | 0.011 |
| IL-2 | NV/NC vs. WCV | 0.0224 | | |
| | WCV vs. RTX | 0.0321 | | |

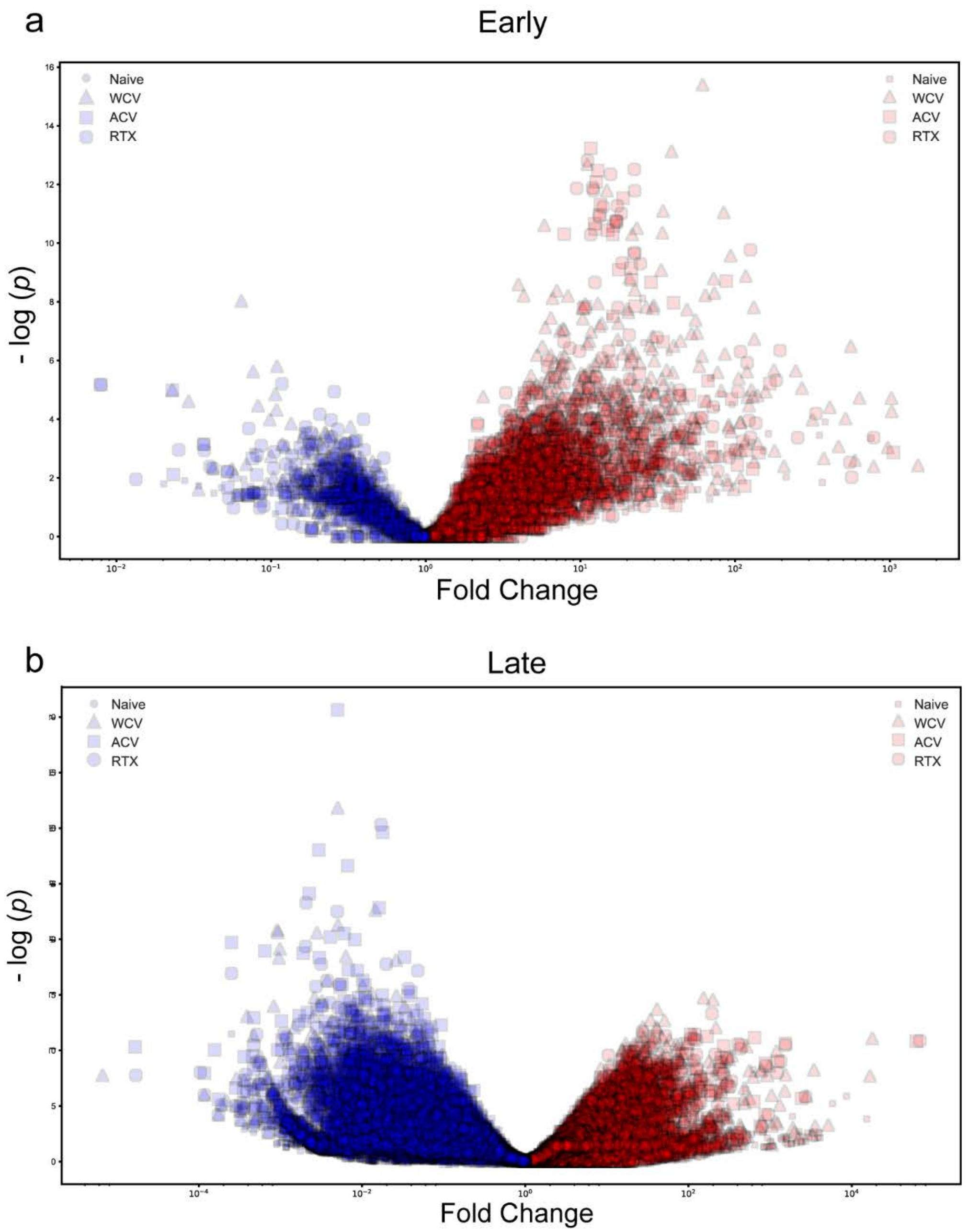


**Supplemental Figure 4: Volcano plot analysis of differentiated genes at early and late time-points of immunized or naïve mice following challenge.** Upregulated genes are represented in red, repressed gens are in blue. Relative fold change was calculated compared to not infected control mice.

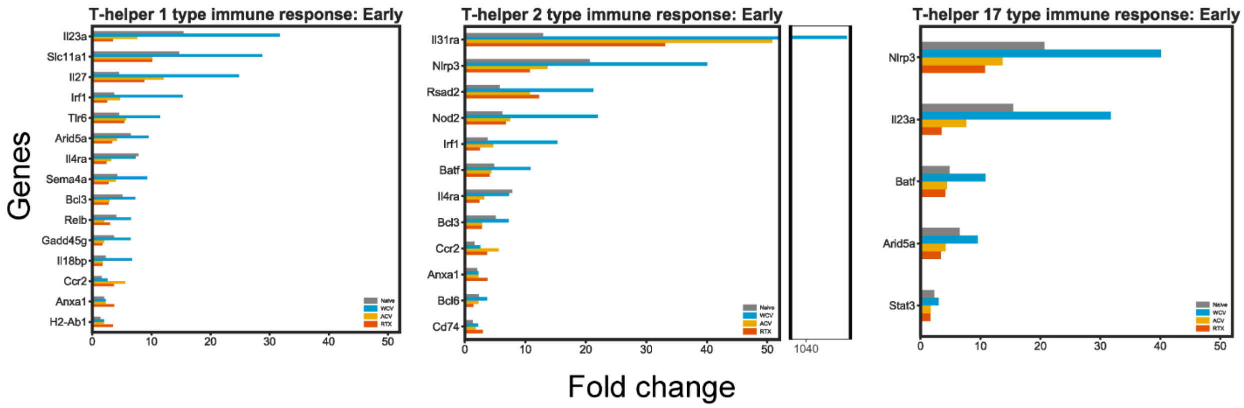


**Supplementary Fig. 5: Gene expression profiles of T helper cell immune responses at early time point.** Differentiated genes associated with (a) Th1, (b) Th2, and (c) Th17 immune responses. Fold changes are relative to NVNC mice. Genes represented are significantly different in at least one group, genes were sorted based on variance of fold change among the experimental groups.

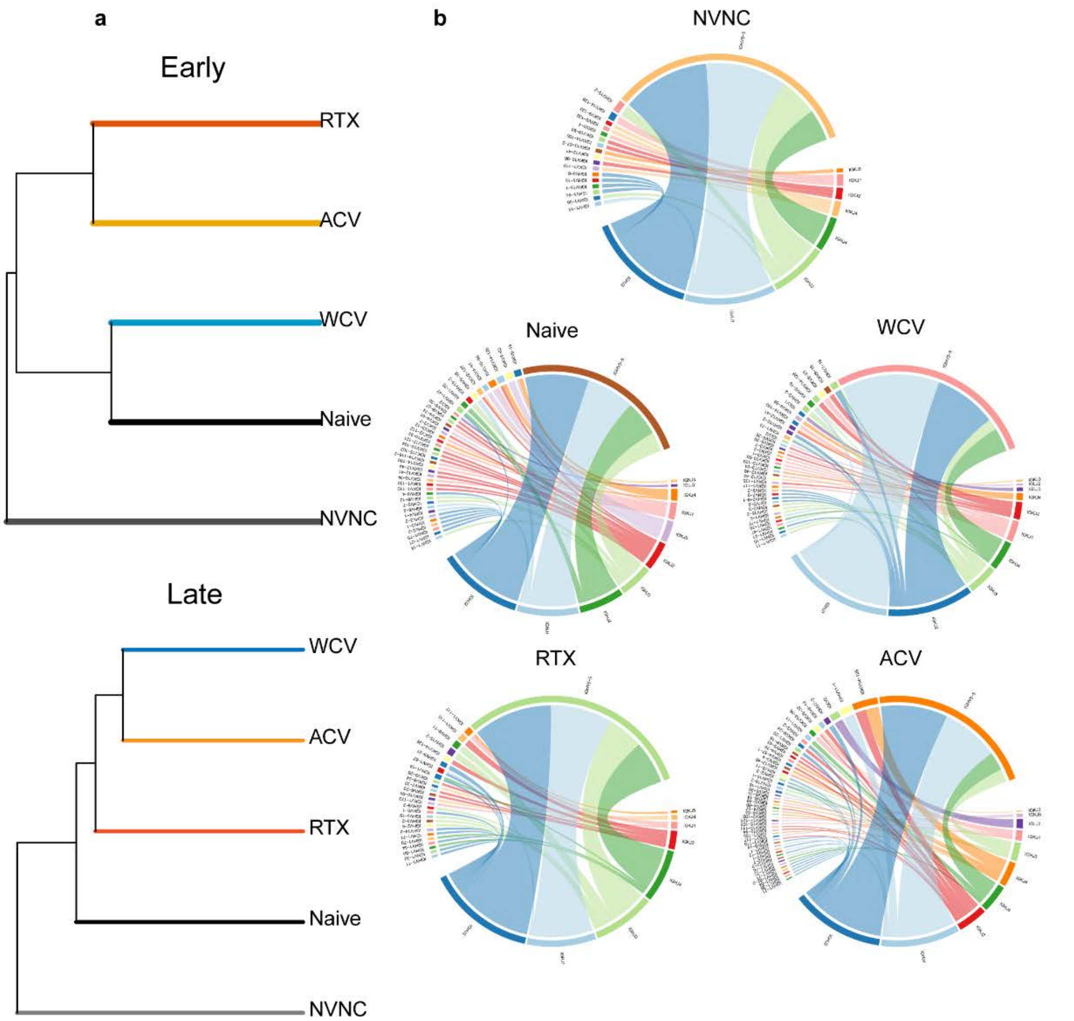


**Supplemental Figure 5:** Repertoire of B cell clones identified from experimental groups at late time point. (a) Relativeness of immunoglobulin profiles between vaccinated and challenged, challenged, or non-challenged groups at late time point. (b) Immunoglobulin diversity and frequency of non-challenged control, experimental groups at late time points. Chord diagrams are used to visualize variable – joining segment diversity. Outer arcs demonstrate overall count of a particular variable (upper portion of chord diagram) or joining segment (lower portion of chord diagram) in each group. Inner ribbons represent the linked segments of a clonotype, while the thickness of the ribbon signifies frequency of the clonotype.
